# Selection for altruistic defense in structured populations

**DOI:** 10.1101/733899

**Authors:** Felix Jordan, Martin Hutzenthaler, Dirk Metzler

## Abstract

We model natural selection for or against an anti-parasite (or anti-predator) defense allele in a host (or prey) population that is structured into many demes. The defense behavior has a fitness cost for the actor compared to non defenders (“cheaters”) in the same deme and locally reduces parasite growth rates. Hutzenthaler et al. (2022) have analytically derived a criterion for fixation or extinction of defenders in the limit of large populations, many demes, weak selection and slow migration. Here, we use both individual-based and diffusion-based simulation approaches to analyze related models. We find that the criterion still leads to accurate predictions for settings with finitely many demes and with various migration patterns.

A key mechanism of providing a benefit of the defense trait is genetic drift due to randomness of reproduction and death events leading to between-deme differences in defense allele frequencies and host population sizes. We discuss an inclusive-fitness interpretation of this mechanism and present *in-silico* evidence that under these conditions a defense trait can be altruistic and still spread in a structured population.

## 1 Introduction

Any trait or character that harms the reproductive success of its bearer while simultaneously increasing the reproductive success of other individuals is called altruistic (e.g., West et al., 2007). The evolutionary success of such traits seems unlikely when considering the individual as the unit of selection (Hamilton, 1963). Nevertheless, numerous examples of altruism are discussed in the literature, such as food sharing in vampire bats (Wilkinson, 1984, 1990), production of costly public goods in bacteria (Diggle et al., 2007; Williams et al., 2007), and even sterility in insects (e.g., Hölldobler and Wilson, 2009). Explanations of how an altruistic trait can evolve are provided by the theories of kin selection (Hamilton, 1964a,b) and group selection (Wynne-Edwards, 1963; Queller, 1992). Also evolutionary game theory has provided important insights into the evolution of altruism and cooperation (Axelrod and Hamilton, 1981; Nowak and May, 1992). The apparent disadvantage of altruism may be overcome if spatial structure or repeated interactions are considered (Nowak, 2006; Ohtsuki et al., 2006).

The theory of kin selection (also known as inclusive-fitness theory) provides the insight that not only the focal individual should be considered when evaluating the success of a trait, but also the effects on their relatives among the interaction partners. The most prominent result of kin selection theory is Hamilton’s rule, stating that an altruistic behavior is selectively favored if the benefit to others, weighted by the relatedness of the actor to the beneficiaries, is greater than the cost to the actor, that is, the reduction of its own offspring (Hamilton, 1964a,b).

An alternative approach to explain the evolution of altruism and cooperation is group selection or multi-level selection theory (see Traulsen and Nowak, 2006; Gardner, 2015, for an overview). When individuals live in distinct groups that are in competition with each other, an advantage on the group level may under certain conditions outweigh a disadvantage on the level of individuals (Wynne-Edwards, 1963; Maynard Smith, 1964; Uyenoyama, 1979; Wade, 1982; Bijma et al., 2007). According to the equation of Price (1970), the selection pressure on a trait is the covariance between the trait and its fitness effects. For the case of subdivided populations, Queller (1992) separated the within- and between-group components of this covariance in Price’s equation. Even if the covariance of a trait with its fitness is negative within each group, it can still be positive in the whole population. This can be understood as an instance of what is known in statistics as the Yule-Simpson effect (Yule, 1903; Simpson, 1951; Blyth, 1972; Sober and Wilson, 1998; Gardner, 2015; Metzler et al., 2016). A necessary condition for this effect is, however, that frequencies of the trait vary between the groups, and an interesting question is what factors besides the kinship structure of the population could maintain this variance over evolutionarily relevant time spans. The theories of kin and group selection have been intensely debated (Traulsen and Nowak, 2006; Nowak et al., 2010; Abbot et al., 2011; Rousset and Lion, 2011; Van Veelen et al., 2012; Gardner, 2015). Birch and Okasha (2015) argue that some of the controversies about inclusive fitness theory and its relationship to group selection theory result from disparate interpretations of the terms in Hamilton’s rule.

Traits of defense against parasites (or predators) can be altruistic as they may be costly for the carrier of the trait while reducing its neighbors’ risk of being infected (or attacked). Costly defense traits are known for many species (Siva-Jothy et al., 2005), including, e.g., resistance against virus infection in meal moths and antibacterial activity in cotton leaf worms (Cotter et al., 2004). Antonovics and Thrall (1994) and Bowers et al. (1994) modeled the dynamics of host–parasite systems for the case that there is an inheritable trait in the host species that makes its carriers less susceptible but also reduces their reproduction rate (see Boots et al., 2009, for a review of extensions of these models). There is, however, no spatial structure in these models and thus no kin or group selection in favor of this resistance trait. Models for the evolution of defense traits in spatially structured host populations have been analyzed by Frank (1998), Brown and Hastings (2003), Schliekelman (2007), Best et al. (2011) and Débarre et al. (2012). Pamminger et al. (2014) analyzed the population structure of ant populations that are the host of a social parasite, a “slavemaking” ant species, and concluded that a defense trait of “slave rebellion” could evolve via kin selection as it reduces the parasite pressure on neighboring ant nests that are likely to be closely related to the “rebels”. Metzler et al. (2016) carried out extensive computer simulation studies for the system of Pamminger et al. (2014) and found that the evolution of slave rebellion may in principle be possible, but only for a narrow range of parameter combinations leading to a meta-population dynamic with kin/group selection on a larger spatio-temporal scale than considered by Pamminger et al. (2014).

Here we analyze further under what conditions a costly defense trait against parasites or preadators can evolve. In the following we will use the terms “host” and “parasite” but our considerations and models can be applied as well to defense against predators in predator–prey systems. Note that in our models we do not assume any kind of adaptive immune system in the host individuals. The defense trait could, however, be an improved innate immunity against the parasite that comes at the cost of a slight fitness reduction and leads to reduced infection rates in the host population.

If defense against a parasite is altruistic and favored by kin or group selection, the success of the costly defense trait depends on the presence of the parasite. Once defenders become abundant, carriers of the defense trait reduce the number of parasites, which in turn may reduce the benefit that each defender provides. It is a priori not clear how this feedback mechanism affects the evolution of the defense trait under kin or group selection. It may for example lead to diminishing returns of the amount of altruism, i.e., the more altruists are present, the lower the additional benefit of even more altruists. Sibly and Curnow (2011) show that coexistence of altruism and cheating will occur if cooperation is subject to diminishing returns. Furthermore, the benefits of defense are mediated by the parasite population, such that there may be a time lag between an altruistic act and the increase in reproductive success of host individuals.

Hutzenthaler et al. (2022) have analytically investigated the scenario of costly defense in structured populations (see also Hutzenthaler and Metzler, 2021), in which carrying the defense trait is imposed with a fitness cost that is independent of the presence of the parasite. For this, Hutzenthaler et al. (2022) used a model of diffusion equations for the frequencies of defenders, cheaters, and parasites and analyzed the asymptotic properties of the model in the limit of weak selection and many demes of large subpopulations with little migration between the demes. For the asymptotic model, Hutzenthaler et al. (2022) have found that defenders will go to fixation if *α*, the fitness disadvantage of defenders compared to cheaters in the same deme, is smaller than the product *β* of the efficacy of defense, the severity of parasites and a birth-death parameter that affects the amount of genetic drift in local host populations. Conversely, defense will go extinct if *α > β* according to the asymptotic model. In Hutzenthaler et al. (2022) the theoretical model analyses were accompanied with simulations that illustrated the basic properties of the model. Here, we perform substantially more extensive simulations to explore the robustness of the asymptotic model predictions of Hutzenthaler et al. (2022).

In Section 2.1 we will first introduce the individual-based model that we use for computer simulations with finite population sizes and finitely many demes to assess the validity of the prediction from the idealized settings in Hutzenthaler et al. (2022). We evaluate the influences of randomness in reproduction, intensity of gene flow, and fluctuations in competition strengths on the success of defense. The asymptotic and heuristic approximations that lead from the individual-based model to the diffusion model of Hutzenthaler et al., 2022, are shown in online appendix A.

Further, we simulate the defender frequency under infinite population sizes and a complete separation of time scales to investigate the effect of these assumptions (sections 2.3). Within this framework we simulate various migration patterns to assess their possible influence on the success of the defense trait. With an extension of the simulation model of Hutzenthaler et al. (2022) we further explore whether the costly defense trait can be considered altruistic or whether it has a long-term direct fitness advantage after several generations (sections 2.4 and 3.3).

## 2 Methods

### 2.1 Model assumptions

Our simulations are based on extensions of the individual-based model of Hutzenthaler et al. (2022) and its diffusion approximation. According to the individual-based model, hosts and parasites populate *D* demes and interact according to a Lotka–Volterra model with within-species competition, combined with a stochastic continuous-time birth– death process. The host population consists of defenders and non-defenders, for short called cheaters in the following, all having the same reproduction rate *g*_*H*_ + *λ*, where *λ* is the growth rate of small host populations in the absence of parasites and *g*_*H*_ is a rate that is added both to the birth and the death rate, modeling random birth and death events that are independent of the host–parasite interactions. Thus, *g*_*H*_ cancels in the average population growth rate but contributes to random genetic drift in the host populations within the demes, that is, random fluctuations of the relative frequencies of defenders.

In addition to the host death rate *g*_*H*_, there is a per-parasite death rate *δ* of each host individual and due to within-species competition a per-host death rate ^*λ*^*/*_*K*_, which implies that *K* is the carrying capacity of the host population in a deme without parasites. Further, we model that defense is a costly trait with an additional death rate *α* of the defenders—independent of the presence of parasites. Thus, if *a*_*i*_ defenders, *c*_*i*_ cheaters and *p*_*i*_ parasites are present in a deme, we obtain

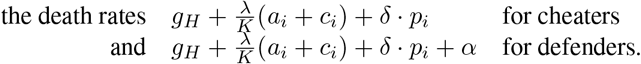

We assume that the parasite population needs the host to grow but also allow for additional random births and deaths with a rate *g*_*P*_. The per-host growth rate of the parasites is *η* but it is reduced by *ρ* for defenders. Thus, the reproduction rate per parasite is *g*_*P*_ + *η c*_*i*_ + (*η* − *ρ*) *a*_*i*_. The parasite death rate is *g*_*P*_ + *ν* + *γ p*_*i*_, where *ν* is the rate at which small parasite populations shrink in the absence of hosts and *γ* is the strength of competition among parasites for other resources than their host.

The migration rates for hosts and parasites are *κ*_*H*_ and *κ*_*P*_. For the individualbased model we assume uniform migration between the demes, that is, the migration rate from any deme *i* to any other deme *j* is *κ*.*/D*. We will consider other migration patterns with the diffusion-model based simulations (see sections 2.3 and 3.2).

To interpret the time scaling, note that if an only-cheater deme population in the absence of parasites reaches its carrying capacity *K*, the death rate equals the reproduction rate *g*_*H*_ + *λ*. The average life span of an individual then is ^1^*/*_(*g*_*H*_ +*λ*)_, and as all individuals reproduce uniformly during their life span, the generation time is roughly ^0.5^*/*_(*g*_*H*_ +*λ*)_.

Taking the limit of many demes, large populations and small values of cost and effects of defense as well as small migration rates and applying some heuristic simplifications (see online appendix A), we can approximate our present model by a model of Hutzenthaler et al. (2022) in which the chance of the defense trait to go to fixation depends on whether the cost *α* of defense is smaller or larger than 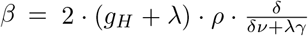. Note that besides the decrease *ρ* in parasite growth rate per defender and the the host death rate *δ* per parasite, which both are clearly relevant for the benefit of the defense trait, *β* contains also the factor *g*_*H*_ + *λ*, which contributes to *genetic drift*, that is, randomness in host reproduction, and thus to between-deme variation, which is needed for deme-level selection or kin selection to be effective.

As the different questions that we address in the following entail different requirements on the computer simulations regarding flexibilty and efficiency, have applied three different simulation approaches: individual-based simulations (section 2.2), diffusion-model based simulations (section 2.3) and simulations updating absolute numbers of individuals after discrete time steps (section 2.4).

### 2.2 Individual-based simulation model for finite population sizes

The initial sizes of defenders, cheaters, and parasites in each deme are *A*_0_, *C*_0_, and *P*_0_. In each simulation step, a single transition is performed, which can be a birth, death, or migration event for one individual of one of the three types in any deme. The inverse of the sum of all transition rates is added to the time that has passed. This corresponds to the expected value of the time until the next transition occurs, which is used instead of the random value for computational efficiency. (The random value is exponentially distributed with a rate of the sum of all transition rates.) The transition is randomly chosen with probabilities proportional to the transition rates. After the numbers of individuals are updated with respect to the randomly chosen transition event, the transition rates are adjusted accordingly. If the time that has passed exceeds the specified time horizon *T* or if defenders or cheaters go to fixation, then the simulation run terminates. Otherwise, the next transition step is performed.

In *simulation series α*, we simulated the individual-based model with various combinations of parameter values for defense disadvantage *α* and for the effect of costly defense, *ρ*, and thus for different values of the parameter 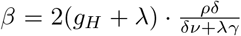 that is crucial for the spread of the defense allele according to Hutzenthaler et al. (2022). To further understand how random genetic drift in the finite model affects the outcome, *simulation series g*_*H*_ was performed for various values of *g*_*H*_.

For finite population sizes *N <* ∞, the time scales of evolutionary and ecological forces are not separated and the fixation probability of the defense trait may depend on the migration rate. A low value of *κ*_*H*_ means that effects within demes dominate the dynamics, while a very large value removes the effect of population structure. We investigated the influence of *κ*_*H*_ in *simulation series κ*_*H*_.

In *simulation series a*_*f*_, we relaxed the assumption of having identical conditions in each deme. After each generation, the competition rate was allowed to change in each deme, independently for hosts and parasites, each with probability *p*_*f*_. If such a change occurred, a normally distributed random number *r* with mean zero and standard deviation ^*af λ*^*/*_*K*_ for hosts or *a*_*f*_ *γ* for parasites was drawn. The new competition rate was then set to *r* + *λK* in the case of hosts or *r* + *γ* in the case of parasites. If the resulting value was non-positive, new random numbers were drawn until a positive value was obtained. The effect of these fluctuations was investigated for different values of *a*_*f*_. The values for all parameters in the different simulation series are shown in Table 1. All simulation results were analyzed using *R* (R Core Team, 2022).

**Table 1:** Parameters of individual-based model with values used in the different simulation series. Entries marked with “var” vary for the simulation series and the values are provided in the corresponding plots.

| Par. | Description | $\alpha$ | $g_H$ | $\kappa_H$ | $a_f$ |
| --- | --- | --- | --- | --- | --- |
| $D$ | number of demes | 50 | 50 | 50 | 50 |
| $T$ | time horizon | $1.25 \cdot 10^6$ | $1.25 \cdot 10^6$ | $1.25 \cdot 10^6$ | $1.25 \cdot 10^6$ |
| $A_0$ | initial defenders per deme | 500 | 500 | 500 | 500 |
| $C_0$ | initial cheaters per deme | 500 | 500 | 500 | 500 |
| $P_0$ | initial parasites per deme | 500 | 500 | 500 | 500 |
| $\lambda$ | host growth rate | 0.5 | 0.5 | 0.5 | 0.75 |
| $K$ | host carrying capacity | 1000 | 1000 | 1000 | 1000 |
| $\delta$ | host death rate per parasite | 0.001 | 0.001 | 0.001 | 0.0005 |
| $\alpha$ | additional death rate of defenders | 0 to 0.01 | 0.001 | 0.001 | 0.001 |
| $\nu$ | parasite death rate | 0.25 | 0.25 | 0.25 | 0.25 |
| $\rho$ | parasite death per defender | 0 to $1.25 \cdot 10^{-4}$ | 0 to $1.75 \cdot 10^{-3}$ | 0 to $1.25 \cdot 10^{-3}$ | 0 to $2.5 \cdot 10^{-3}$ |
| $\gamma$ | parasite competition | 0.002 | 0.002 | 0.002 | 0.0015 |
| $\eta$ | parasite growth per host | 0.001 | 0.002 | 0.002 | 0.004 |
| $\kappa_H$ | host migration rate | 0.001 | 0.001 | $10^{-5}$ to 0.01 | 0.001 |
| $\kappa_P$ | parasite migration rate | 0.001 | 0.001 | 0.001 | 0.001 |
| $g_H$ | randomness in host reprod. | 0.5 | 0 to 5 | 0.5 | 0.5 |
| $g_P$ | randomness in parasite reprod. | 0.5 | 0.5 | 0.5 | 0.5 |
| $p_f$ | probability of fluctuating compet. | 0 | 0 | 0 | 0.02 |
| $a_f$ | amount of compet. fluctuation | 0 | 0 | 0 | 0 to 0.25 |

### 2.3 Simulation model for defender frequency in large populations

In the simulation series to check possible effects of different population structures we applied a simulation approach that is based on the diffusion approximation *X*_*t*_ = (*X*_*t*_(1), …, *X*_*t*_(*D*)) for the relative frequencies *X*_*t*_(*i*) of defenders in all demes *i* ∈ 𝒟 := 1, …, *D* at time *t*, as derived by Hutzenthaler et al. (2022) for the limit of large populations, weak selection and little migration, see also online appendix A.

In Hutzenthaler et al. (2022), the defenders frequency on the evolutionary time scale (i.e., speeding up time by a factor of *N*) in the limit of large population sizes (scaled with the same factor *N* → ∞) is shown to be described by the diffusion equations

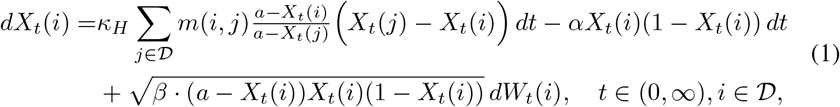

where {*W* (*i*): *i* ∈ 𝒟} are independent standard Brownian motions. This defines a process *X*_*t*_. To simulate this process we chose a small, positive step size *dt* (of 10^−5^ time units), and in each simulation step corresponding to a time interval [*t, t* + *dt*] and all *i* ∈ {1, …, *D*} we simulated *X*_*t*+*dt*_(*i*) = *X*_*t*_(*i*) + Δ_*t*_(*i*), where Δ_*t*_(*i*) is normally distributed with

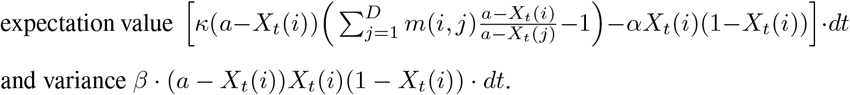

With 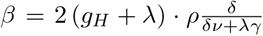 and 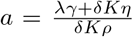 this approximates the dynamics of defense allele frequencies in the model of section 2.1 (see online appendix A). The migration rate *m*(*i, j*) from deme *j* to deme *i* depends on the migration scheme, see below.

At the beginning of the simulation, *X*_0_(*i*) is set to *x*_0_ in each deme *i* ∈ {1, …, *D*}. If *X*_*t*_(*i*) *< ε*, then *X*_*t*_(*i*) is set to zero, or if *X*_*t*_(*i*) *>* 1 − *ε*, then *X*_*t*_(*i*) is set to one. In each step, *t* is increased by *dt*, and if *t* is an integer-valued multiple of ^*T*^*/*_1000_, then the average value of *X* across all demes is computed. If this average value is zero or one, or if *t* ≥ *T*, then the simulation run is terminated.

#### 2.3.1 Migration models and parameter ranges

To assess how well the results from the theoretical analysis could predict the outcome for finitely many demes, we performed simulations for a fixed value of defense disadvantage *α* and different values of the benefit *β*. This was done for different values of the number of demes, *D*, to check how well the setting could be approximated by the assumption of infinitely many demes in the theoretical analysis. In further analyses, the robustness of the system towards various migration schemes was investigated, which were as follows, where *m*(*i, j*) refers to migration from deme *j* to deme *i*:

**U:** Uniform migration between all demes. *m*(*i, j*) = ^1^*/*_*D*_ for all *i, j* ∈ {1, …, *D*}

**N:** Stepping-stone model on a one-dimensional series of demes, closed to a circle. *m*(*i, j*) = ^1^*/*_2_ if |*i* − *j*| = 1 or {*i, j*} = {1, *D*}; *m*(*i, j*) = 0, otherwise

**2D:** Stepping-stone model on a two-dimensional lattice of demes on a torus, that is, migration only on to the four neighboring demes *m*(*i, j*) = ^1^*/*_4_ if *i* next to *j* on two-dim. lattice on a torus; *m*(*i, j*) = 0, otherwise

**T:** The demes are the nodes of a binary tree; migration only along the branches of the tree. *m*(*i, j*) = ^1^*/*_3_ if *j >* 1 is parent or child of *i* or if *i* and *j* are neighboring leaves; 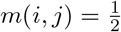, if *j* = 1 and *i* ∈ {2, 3}; *m*(*i, j*) = 0, otherwise

The parameters used in the different settings are shown in Table 2. For each combination parameter values we performed 100 simulations.

**Table 2:** Parameters of diffusion simulations with values for runs with uniform migration (U), nearest-neighbor migration (N), two-dimensional nearest-neighbor migration (2D), and migration along edges of a binary tree (T). Entries marked with “var” vary for the simulation series and the values are provided in the corresponding plots.

| Par. | Description | U | N | 2D | T |
| --- | --- | --- | --- | --- | --- |
| $\beta$ | random fluctuation (“benefit”) | 0.01 to 0.1 | 0.01 to 0.1 | 0.02 to 0.18 | 0.01 to 0.1 |
| $\alpha$ | defense disadvantage | 0.05 | 0.05 | 0.1 | 0.05 |
| $\kappa$ | migration rate | 0.1 | 0.1 | 1 | 0.1 |
| $a$ | see Section 2.3 | 2 | 2 | 2 | 3 |
| $D$ | number of demes | 5 to 500 | 5 to 500 | 9 to 484 | 7 to 511 |
| $x_0$ | initial defense frequency per deme | 0.5 | 0.5 | 0.5 | 0.5 |
| $T$ | time horizon | 2000 | 2000 | 2000 | 2000 |
| $dt$ | time step size of discretization | $10^{-5}$ | $10^{-5}$ | $10^{-5}$ | $10^{-5}$ |
| $\varepsilon$ | cutoff value | $10^{-7}$ | $10^{-7}$ | $10^{-7}$ | $10^{-7}$ |

### 2.4 Simulation to assess direct long-term fitness and long-term altruism

Being a defender in our model is costly in the sense that defenders produce offspring at a lower rate than non-defenders in the same deme. This alone does not immediately imply that defense is costly in terms of progeny numbers after several generations. One could imagine that by reducing the parasite load, defense behavior of some host individual could lead to more progeny after a few generations than in the case that the individual would not defend but still live in the same environment. In this case, defense behavior would not be costly in terms of long-term direct fitness, that is number of progeny after several generations. To assess the long-term fitness costs and benefits of defense behavior we carried out simulations in which a random defender was chosen and became a *traitor*, that is, an individual that does not behave as a defender and does not pay the costs of defense but still has the defense allele and passes it on to their offspring. We call the defense behavior *long-term costly* (and say that treason has a long-term direct fitness advantage) if from a certain number of generations on the expectation value for the number of progeny is higher in the case in which the defender becomes a traitor. In the evaluation of our simulation results we choose the more practical approach to assess after a decent number of generations (≈50) how many progeny the traitor had compard to simulation in which the chosen defender behaved as a defender. If a behavior is long-term costly and beneficial for other host individuals (e.g., in the same deme), we also call it *long-term altruistic*.

Our simulation model is based on the finite-population model specified in section 2.1 with 200 demes and equal migration rates between all pairs of demes. Like in Hutzenthaler et al. (2022) we applied the *τ* -leaping approach in the simulation program (Gillespie, 2001), which is faster than the individual-based simulations in section 2.2 For this, we calculated for short time spans *τ* for each type of event, e.g., reproduction of some of the *n* defenders in a particular deme, the per-individual rate *r* (in the example *r* = *g*_*H*_ + *λ*) of this event and drew a (*n, r* · *τ*)-binomially distributed number of individuals for whom this event takes place. The time step length *τ* was chosen small enough to prevent that *r* · *τ* became larger than 0.01 (at least if the host population is not larger than its carrying capacity in the absence of predators).

Our simulation runs started with initializing each deme with random numbers of defenders, cheaters and predators. Each simulation run began with a first phase (called phase 1 in the following), in which we simulated 20 million time steps, corresponding to approximately 400,000 host generations. If defenders were not extinct after phase 1, we continued with phase 2, which consisted of 1000 iterations. In each of these iterations we purely-randomly chose a defender and continued with two different simulations, one in which the chosen individual was turned into a traitor and one without traitors. Note that in each such iteration, at most one individual could become a traitor, and the offspring of traitors were defenders. The two simulations (with the chosen indivdual as traitor or as normal defender) were coupled by using the same series of pseudo-random numbers as the basis for (pseudo-)binomially distributed numbers. This means that if in the exact same step of the two simulations the parameters of the two binomial distributions were identical (e.g., as that deme at that time was by no means affected by the chosen individuals or their offspring), the two binomial pseudorandom variables gave the same value. Also when the two corresponding binomial random variables from the simulations with and without traitors had different parameters, they were still coupled, as in this case we generated for both simulations first the same value *U* from a uniform distribution between 0 and 1 and then converted this value into binomial pseudo-random values by applying the inverse of the distribution function of each binomial distribution to the value *U*. We then compared the progeny numbers of the chosen individual in the two simulations after 25 simulated time units (that is, approximately 50 generations). This procedure of phase 1 and a possible phase 2 with 1000 iterations was repeated for various parameter combinations and also with replications including independent simulations of phase 1.

## 3 Results

### 3.1 Finite populations

In simulation series *α, g*_*H*_, *κ*_*H*_, and *a*_*f*_, the number of demes was set to *D* = 50 and the initial numbers of defenders, cheaters and parasites per deme were 500 each. For the values of the parameters, see Table 1. For each parameter configuration, 100 simulation runs were performed. In *simulation series α* we tested the effect of finite population sizes on the theoretical prediction of Hutzenthaler et al. (2022) for different values of *α* and *β*. The mean values of the results of the 100 simulations are shown in Figure 1a. All final frequencies were either zero or one, i.e., extinction or fixation of defense was observed in each run, such that the final frequencies in Figure 1a are the fractions in which the defense type went to fixation. The results agree remarkably well with the prediction of the asymptotic model that defenders have a higher fixation probability than cheaters for *α > β* and vice versa for *α < β* (thresholds are shown as dashed lines). But in contrast to the predictions of the asymptotic model, it is not certain which of the two types will go to fixation if *β* is in a certain range around *α*.

**Figure 1:**
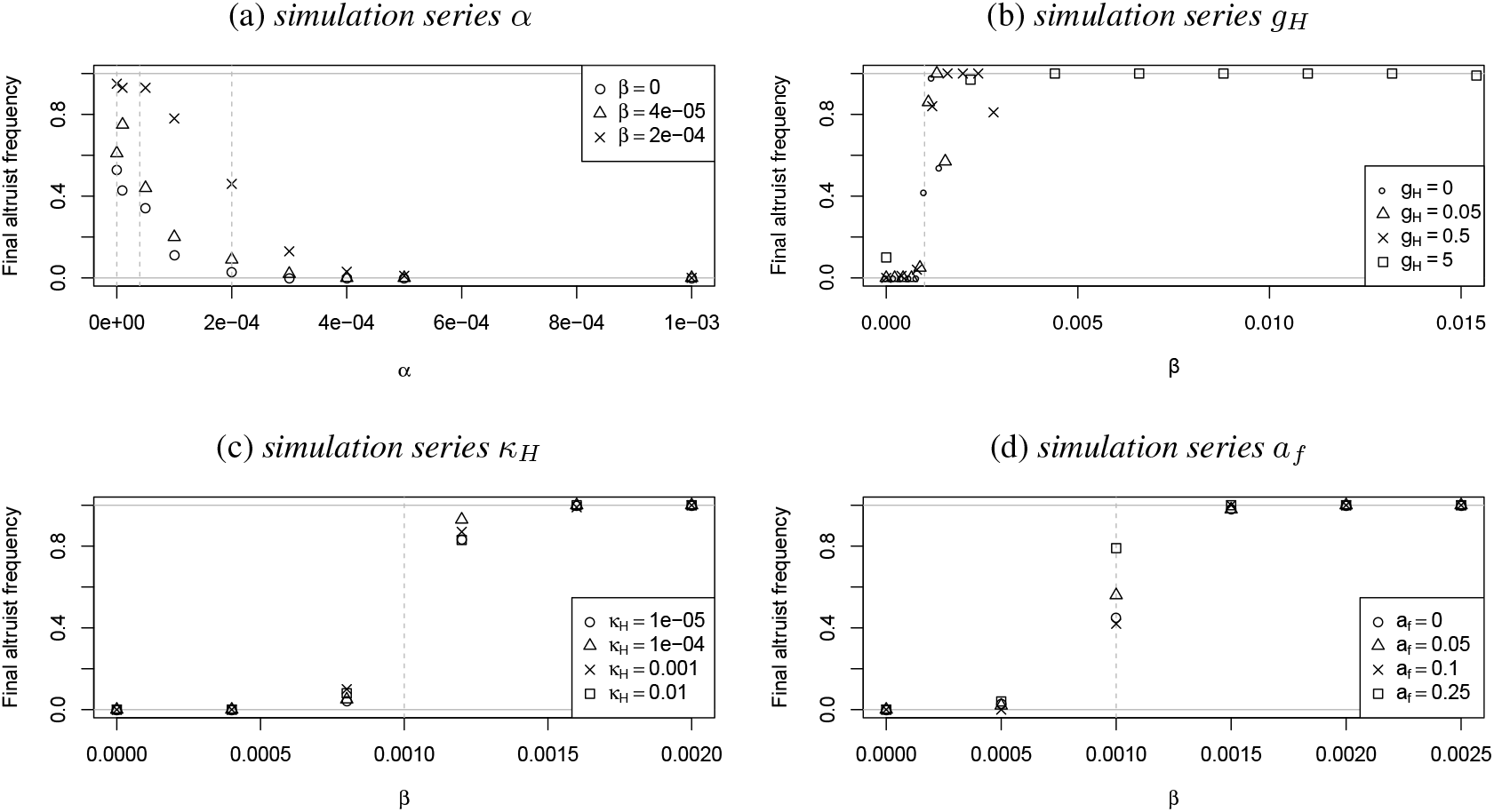
Average final defender frequency across 100 runs, which is in most cases the fraction of simulations in which the defense trait went to fixation. Dashed lines show threshold for defense advantage according to asymptotic model. See Section 3.1 for details.

For *α* = 10^−3^, extinction of defense was observed in all 100 simulations for each of the values of *β*. For *α* = 5 · 10^−4^ and *α* = 4 · 10^−4^, there was no significant difference between the results for the different values of *β* (Bonferroni-Holm corrected p-values of 0.37 and 0.10, respectively). For all other values of *α*, there was significant difference between the results for the different values of *β* (all Bonferroni-Holm corrected p-values below 0.0002).

Effects of randomness in host reproduction introduced by *g*_*H*_ were investigated in *simulation series g*_*H*_ (see Figure 1b). Note that the different *β* values in this simulation series resulted from combining the values for *g*_*H*_ with the *ρ* values 0, 2.5 · 10^−4^, 5 · 10^−4^, 7.5 · 10^−4^, 1 · 10^−3^, 1.25 · 10^−3^, 1.5 · 10^−3^ and 1.75 · 10^−3^. There was no significant difference between the results for the different values of *g*_*H*_ for *ρ* = 1.5 · 10^−3^ (Bonferroni-Holm corrected p-value of 0.11). For all other values of *β*, there was a significant difference (maximum corrected p-value of 2 · 10^−6^). In the simulations with *ρ* = 1.75 · 10^−3^, the parasite population went extinct in 94, 98, 96, and 82 runs out of the 100 runs for *g*_*H*_ = 0, *g*_*H*_ = 0.2, *g*_*H*_ = 2, and *g*_*H*_ = 20, respectively. For all other values of *β*, the parasite population never went extinct in any of the simulations.

In *simulation series κ*_*H*_, we investigated the effect of host migration rates. Results are shown in Figure 1c. Among the simulations with *κ*_*H*_ = 10^−5^, there were eight and five runs out of 100 total, for *β* = 8 · 10^−4^ and *β* = 1.2 · 10^−3^, respectively, where neither extinction nor fixation of defense had occurred by the end of the simulation. This means there were both defenders and cheaters present in the host population and final defenders frequency was strictly between zero and one. In all other simulations of series *κ*_*H*_, either the defenders or the cheaters went extinct. In the case of *β* = 8 · 10^−4^, the defenders went extinct in 88, 95, 90 and 92 of 100 simulations with the different values of *κ*_*H*_, and for *β* = 8 · 10^−4^ they went to fixation in 79, 93, 87 and 83 of the the 100 simulations. For *β* = 0 and *β* = 4 · 10^−4^, all simulations led to extinction of defenders, while for *β* = 1.6 · 10^−3^ and *β* = 2 · 10^−3^, all simulations showed fixation of defense. Intermediate values of *β* showed no significant difference between different settings of *κ*_*H*_ (all Bonferroni-Holm corrected p-values above 0.39).

The effect of random fluctuations in host carrying capacity was studied in *simulation series a*_*f*_. Results are shown in Figure 1d. For *β* = 0, all simulations led to extinction of defenders, while for *β* = 2 · 10^−3^ and *β* = 2.5 · 10^−3^, all simulations showed fixation of the defense trait. There was a significant effect of the amount of fluctuation, *a*_*f*_, for *β* = 0.001 (Bonferroni-Holm corrected p-value: 5.6 · 10^−7^). For *β* = 5 · 10^−4^ and *β* = 0.0015, the effect of *a*_*f*_ was not significant (corrected p-value of ≈ 0.52 for both).

In most simulations with the finite-population models the defense trait either went to fixation or to extinction. When *α < β* it did not always go to fixation and when *α > β* it did not always go extinct as would be the prediction of the asymptotic model. However, in all four finite-population simulation series, extinction of the defense trait was more frequent than fixation for the parameter combinations with *α > β* and fixation of the defense trait was more frequent than extinction for all parameter combinations with *α < β* (Figure 1d).

### 3.2 Large populations

Several simulation series were performed to check how well the condition derived in the theoretical analysis could predict the outcome in a setting of large populations living in finitely many demes with different migration patterns. The parameters of all simulation series are shown in Table 2. First, *simulation series ε* was performed to find a suitable cut-off value for the discretization (see online appendix B for details), resulting in *ε* = 10^−7^.

The value *ε* = 10^−7^ was then used in *simulation series U, N, 2D*, and *T*, with 100 runs per combination of parameter values. Mean values with standard errors from these simulation series are shown in Figure 2, and root mean squared errors of the simulation results compared to the theoretical prediction are shown in online appendix C in Figure 2.

**Figure 2:**
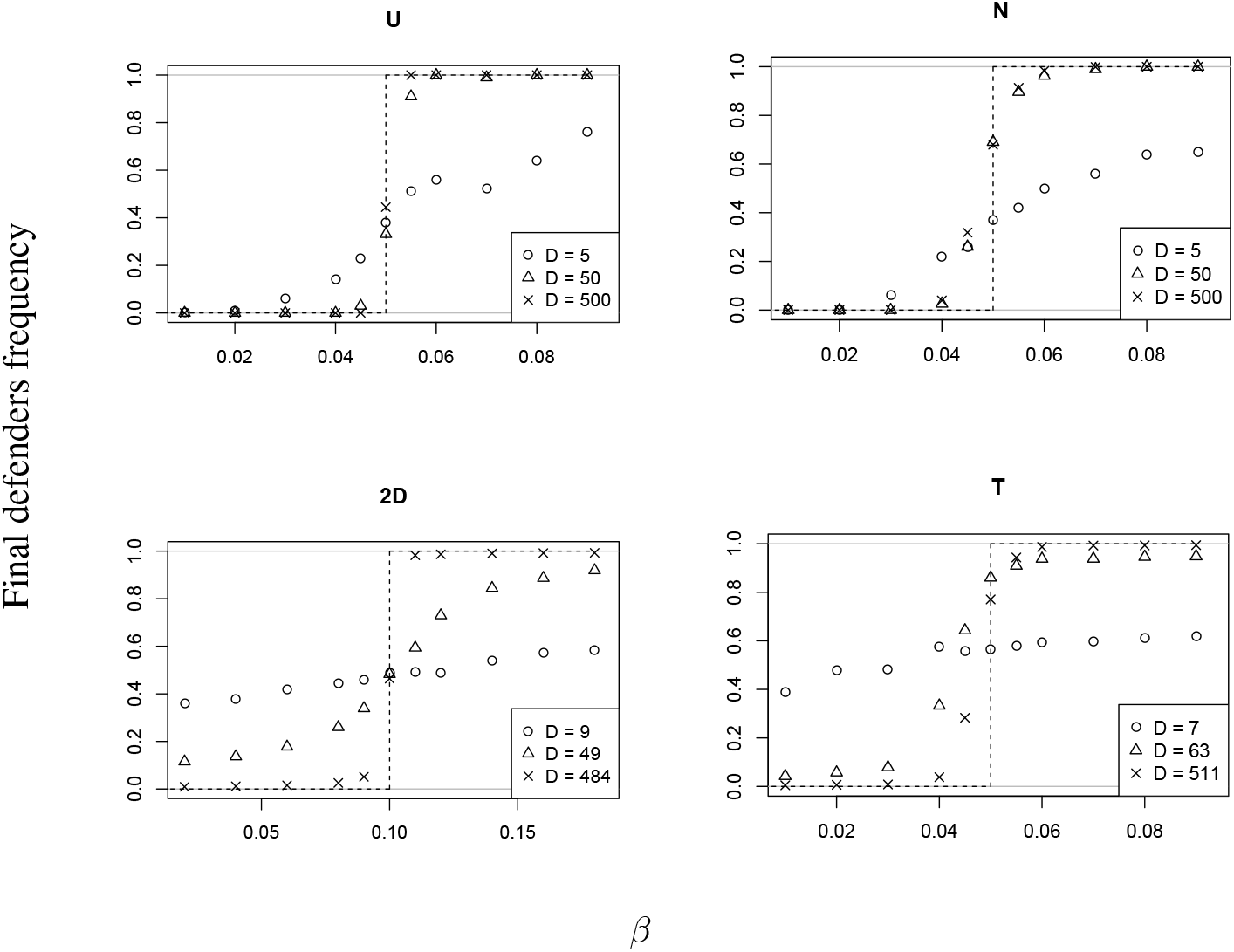
Mean values over 100 simulation runs with uniform migration (U), nearestneighbor migration (N), two-dimensional nearest-neighbor migration (2D), and migration along edges of a binary tree (T). Dashed lines indicate the theoretical predictions.

In *simulation series U* we checked the prediction from the theoretical results for uniform migration between the demes, with different total deme numbers. For each parameter setting, 100 simulation runs were performed. Among the simulations with *β* = *α*, 84 runs with *D* = 500 and one run with *D* = 50 showed neither extinction nor fixation of the defense trait. All other runs had a final defenders frequency of either zero or one (see Figure 3 in online appendix C for details).

**Figure 3:**
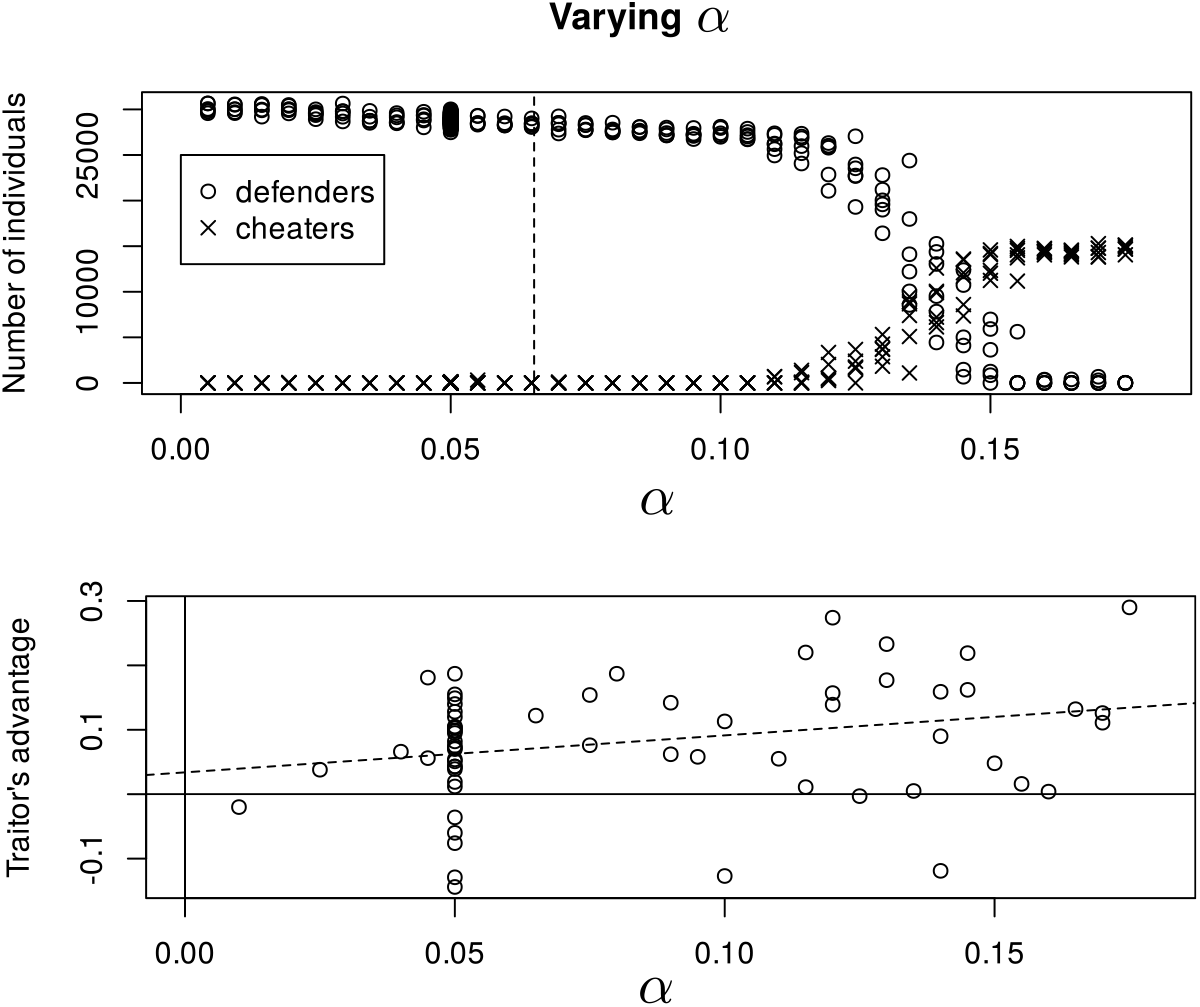
Simulation results with various values of the defense cost *α* and default values of section 3.3 for the other parameters. Top: total numbers of defenders and cheaters after phase 1 (see section 2.4) and asymptotic threshold (dashed line). Bottom: Scaled difference of ultimate progeny numbers of a defender when becoming a traitor or not, and the dashed line is a weighted-regression line; only non-zero values are shown. See section 3.3 for further details.

In *simulation series N* we simulated the process *X* under nearest-neighbor migration on a circle. For each parameter setting, 100 simulation runs were performed. For *D* = 50, 125 of the 500 runs with 0.04 ≤ *β* ≤ 0.06 showed coexistence of defenders and cheaters, while for *D* = 500, coexistence was observed in 425 of the 700 runs with 0.04 ≤ *β* ≤ 0.08. All other runs of the simulation series showed fixation or extinction of defenders. For *α* = *β*, a final defenders frequency of either zero or one is observed in 100, 45, and 0 runs (out of a total of 100 runs) for *D* = 5, *D* = 50, and *D* = 500, respectively (See Figure 4 in online appendix C).

**Figure 4:**
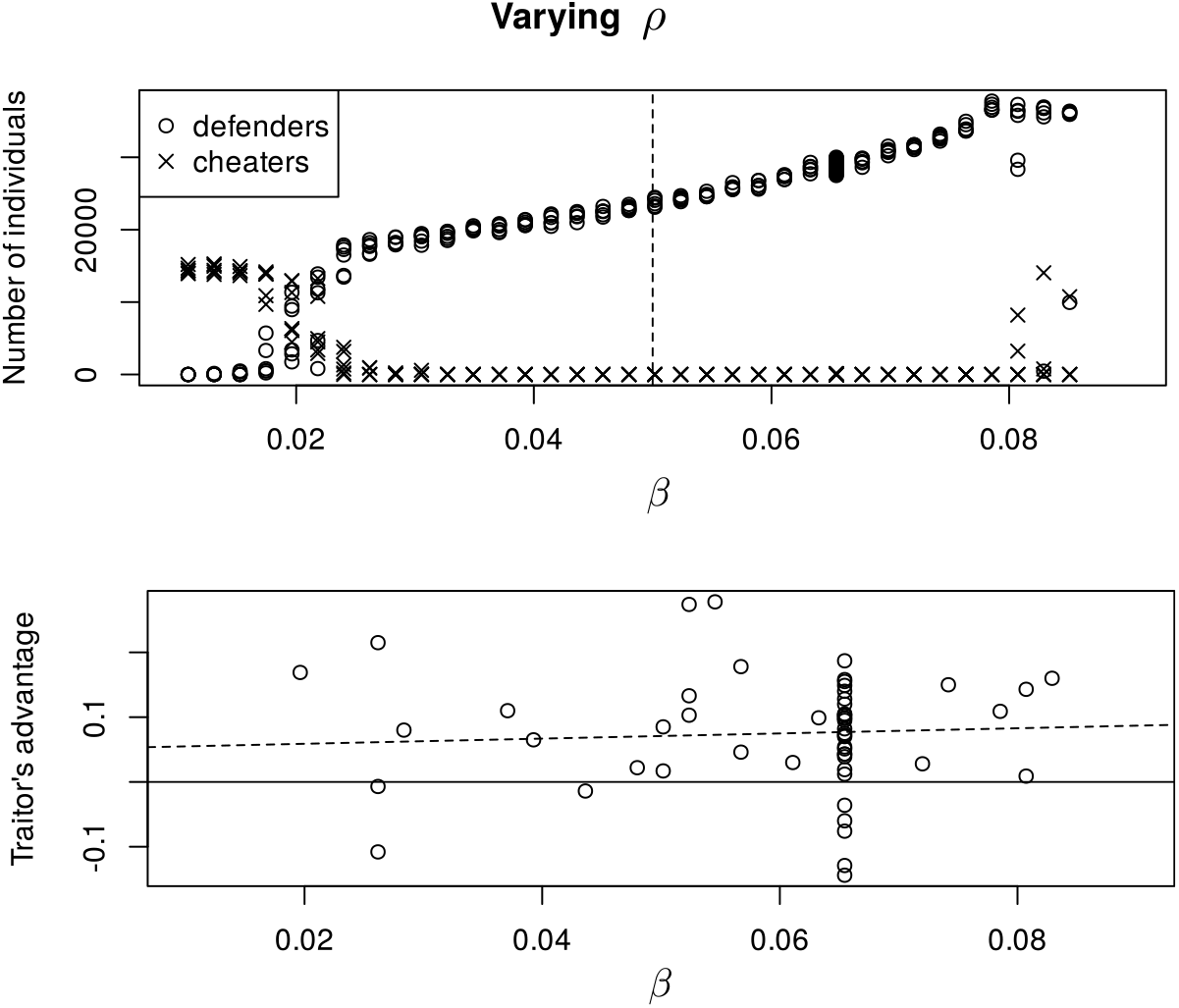
Results of simulations with various values of the defense efficacy *ρ*, resulting in different values of *β*. See caption of Fig. 3 and section 3.3 for details.

In *simulation series 2D* we assumed nearest-neighbor migration in two dimensions, i.e., each deme sent migrants to its four immediate neighbors (using a torus structure to avoid boundary effects). For each parameter setting, 100 simulation runs were performed. In none of the simulations, fixation or extinction of defense occurred, but with *D* = 22^2^ = 484, the final defenders frequency was always close to 0 if ^*β*^*/*_*α*_ *<* 0.8 and close to 1 if ^*β*^*/*_*α*_ *>* 1.1 (Figure 5 of online appendix C).

**Figure 5:**
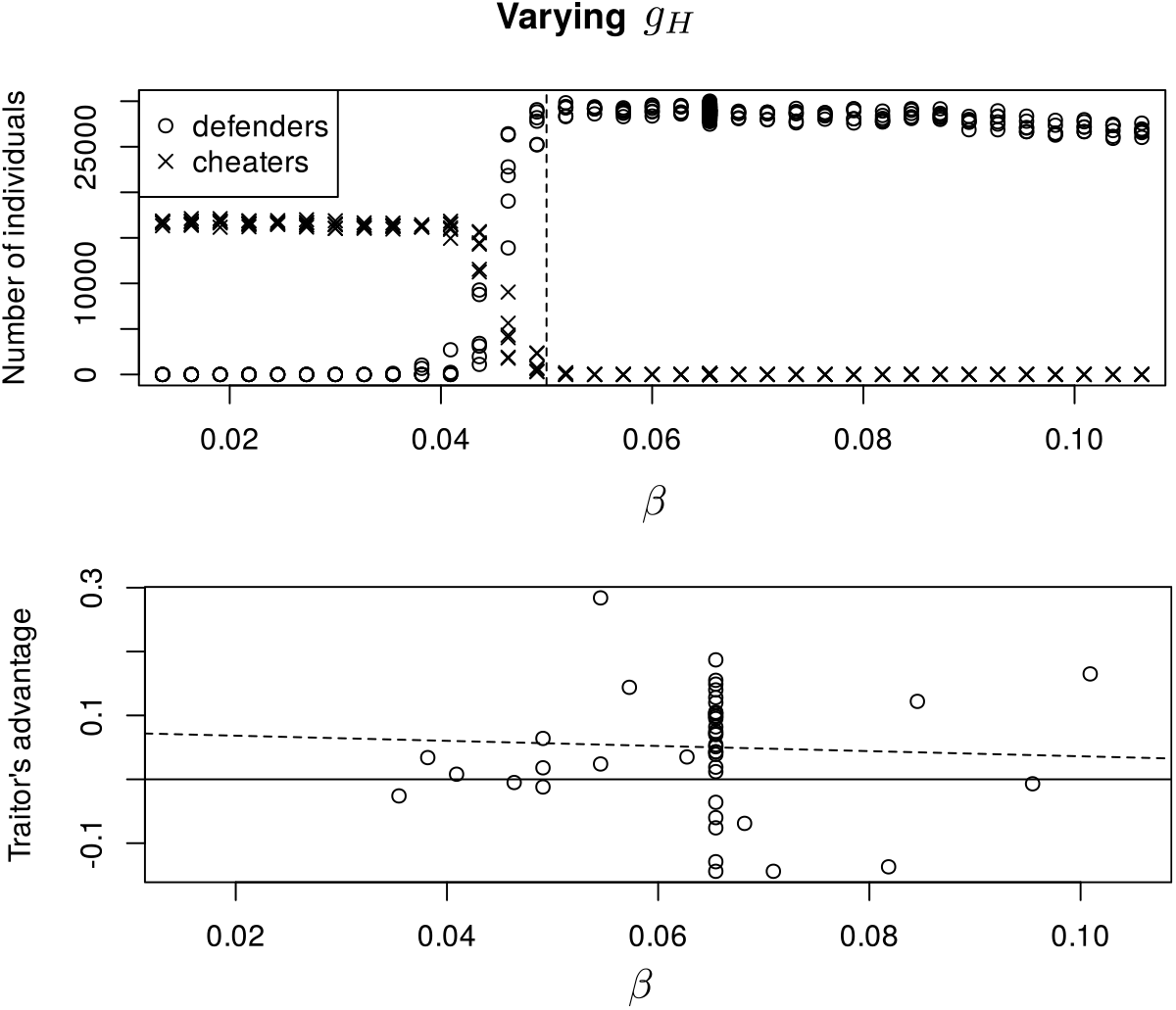
Results of simulations with various values of the parasite-independent host birth–death rate *g*_*H*_, affecting genetic drift and resulting in different values of *β*. See caption of Fig. 3 and section 3.3 for details.

We simulated the process *X* with migration along edges of a binary tree in *simulation series T*. Demes were represented as nodes of a binary tree and each node exchanged migrants with its parent and its two children. The root sent migrants only to its two children. Neighboring leaves also exchanged migrants. For each parameter setting, 100 simulation runs were performed. In all simulations, the final defenders frequency was strictly between 0 and 1, but for *D* = 511 very close to 0 if ^*β*^*/*_*α*_ *<* ^0.04^*/*_0.05_ = 0.8 and very close to 1 if ^*β*^*/*_*α*_ *>* ^0.06^*/*_0.05_ = 1.2 (Figure 6 in online appendix C).

### 3.3 Long-term fitness and long-term altruism

To explore the direct long-term fitness of defense, we simulated data according to the model specified in section 2.4 with the parameter values *λ* = 1, *g*_*H*_ = 5, *K* = 200, *δ* = 0.002, *α* = 0.05, *g*_*P*_ = 0, *η* = 0.04, *ρ* = 0.03, *ν* = 0.5, *γ* = 0.01, *κ* = 0.001 and *ι* = 10^−5^, which we call *default parameters values* for the rest of this section. Note that this combination of parameter values is further away from the large-populations asymptotics than the parameter values were in the finite-population simulations in section 3.1, e.g., with smaller host populations in the demes (host carrying capacity *K* = 200). Our main reason for choosing this parameter combination for this model was computational efficiency as it allowed us to obtain in acceptable time a decent number of simulations in which changing a defender into a traitor had an effect on its long-term fitness. As an additional benefit, using these parameter values allowed us also to further explore the range for which the asymptotic results are applicable. To check, however, whether the two simulation approaches (sections 2.1 and 3.1 compared to section 2.4) lead to similar results when applied with the same parameter values, we carried out some simulations with the simulation program from 2.4 with parameter values as in some of the simulations in section 3.1 and found that the results are in accordance to the predictions of the asymptotic model as much as the results in section 3.1, see online appendix D.

We launched 240 independent simulation runs with the default parameter values. (Here and in the following we refer with “simulation run” to a simulation consisting of phase 1 of the initial 20 million time steps, corresponding to approx. 400,000 host generations, and phase 2, in which 1000 defenders were chosen and for each of them their descendants for approximately 50 generations were simulated with and without the chosen defender being a traitor.) In 26 of these runs there was exactly one of the 1000 chosen defenders for which the final progeny number was different in the traitor simulation than in the non-traitor simulation, and for two of the 240 runs there were two chosen defeders with this property. For each of these 28 runs we calculated a measure for the average advantage of treason by dividing the difference between the number of traitor progeny and the corresponding number without treason by 1000 (the number of chosen pairs). Further we weighted this number proportionally to the number of defenders after phase 1 of the simulation run, with the average weight being 1. The mean value of these 28 weights was 0.0571 and significantly larger than 0 (*p* ≈ 0.0011, two-sided t-test; Studentized 95% confidence interval [0.0251, 0.0892]).

For each of the model parameters *α* (defense cost), *ρ* (effect of defense) and *g*_*H*_ (randomness in host reproduction) we launched 240 additional simulation runs varying this parameter (and with default values for all other parameters). The results of these three simulation results are shown in Figures 3, 4 and 5, always combined with the results from the 240 runs with default parameters. In their upper plots these figures show the numbers of defenders and cheaters in each of the simulation runs at the end of simulation phase 1. The dashed line shows value of *β* ≈ 0.0654 in the case of Fig. 3 and of *α* = 0.05 of Figs. 4 and 5, and thus the values that are according to the predictions of the asymptotic model the thresholds for the value on the horizontal axis for defense to have an (inclusive and/or multi-level) fitness advantage. In contrast to the results with the parameter ranges in online appendix D, the simulation results in Figures 3, 4 and to a lesser extent also 5 show that the parameter range in which the defense allele can spread is noticeably different from the predictions. Fig. 3 shows that defense has a fitness advantage for *α <* 0.13 (and not only for *α < β* ≈ 0.0654) and Fig. 4 shows that defense was advantageous for *ρ* values that result in *β >* 0.02 (and not only *β > α* ≈ 0.05).

The bottom plots in Figs. 3, 4 and 5 show the traitors’ advantage according to phase 2 of the simulations. If for one of the 1000 chosen defenders of a simulation run the number of progeny at the end of the simulation was different between the two simulations, a data point shows the difference (traitor progeny number minus defender progeny number) divided by 1000, that is scaled per number of simulated individuals. For the dashed lines the mean of these values were taken for each simulation run and a weighted linear regression was applied with weight being the number of defenders individuals after phase 1. Within the simulated parameter ranges we could not identify a parameter combination for which treason appeared to be disadvantageous in terms of long-term direct fitness.

## 4 Discussion

Hutzenthaler et al. (2022) found a criterion to determine the success of a costly defense allele for deme-structured populations in the limit of large populations in many demes under a complete separation of time scales. Here we found for several different simulation models with finite populations or different scenarios of population structure, that this criterion can still be applied to predict whether the defense trait will rather go to fixation or to extinction. The prediction is that the defense allele will rather go to fixation than to extinction if and only if its direct fitness cost *α* is smaller than a parameter *β* (see section 2.1). In the asymptotic diffusion model of Hutzenthaler et al. (2022) for the relative frequencies of the defense trait in each deme, *β* is a factor in the diffusion term. In terms of population genetics, the random genetic drift acting on the relative frequency of the defense allele in a deme is proportional to 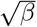. An interpretation from the perspective of multi-level selection is that this genetic drift leads to deme-level variation that is needed for deme-level selection effects. Demes with a high defenders frequency are better protected against parasites and thus have a higher host density. Thereby, they are able to send off more migrants than demes with few defenders. Thus, even though being individually disfavored in each deme, defenders can have an advantage over cheaters in the total population.

### 4.1 Finite-size populations

To approximate the finite-population models by the asymptotic diffusion model, Hutzenthaler et al. (2022) defined *β* by

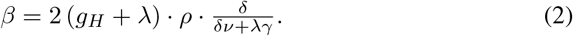

With this, the prediction of the diffusion model that defenders become more frequent than defectors if and only if *β > α* fits very well to our simulation results with a range of different finite population-size models and parameter ranges as shown in table 1 (see section 3.1 and online appendix D). We find the precision of this fit quite remarkable as equation (2) is not only based on rigorous asymptotics but also on the heuristic approximation shown in online appendix A.

The parameter *β*, or at least the product of the last two factors in the right-hand side of equation (2), can be interpreted as the benefit of defense, as *ρ* is the effect that each defender has against the parasite and 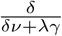 is a scaling of the potential harm *δ* that hosts suffer from parasites. The factor *g*_*H*_ + *λ* is the host reproduction rate. When the system is in a quasi-equilibirium on the ecological time scale, the death rate is equal to the birth rate and *g*_*H*_ + *λ* has almost no effect on expectation values for short-time population size changes, but genetic drift is proportional to 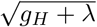, which is important for between-deme variation also in the finite-population model. The diffusion approximation shows that the genetic drift arising from *g*_*H*_ + *λ* is enhanced by the factors *ρ* and 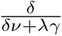 as the genetic drift of the defense allele frequency in the asymptotic diffusion model is proportional to the square root of the product of all three factors.

The simulation results in section 3.1 are based on parameter values that lead to smaller deme-population sizes, as the carrying capacity of each deme for the host population was only 200, even in the absence of the parasite. In this case the parameter range for which defender could become more frequent than cheaters was larger than the predicted by the *α < β* criterion. For this it might play a role that smaller population sizes allow for a metapopulation dynamic with local extinction and recolonization of demes.

### 4.2 Effect of population structure

Simulations assuming infinite population sizes were performed in order to compare the success of defenders in finite and infinite settings and to investigate effects of population structure. We simulated the process *X* given by equation (1). It describes the relative frequencies of defenders in the limit of large population sizes under a complete separation of time scales.

The simulations with uniform migration among 500 demes fit the theoretical prediction very well (Figure 2). The analytical result could only be obtained in Hutzenthaler et al. (2022) when uniform migration is assumed, allowing for a mean field approximation. Nevertheless, for the other migration schemes investigated, the prediction was also closely met, when the number of demes was large enough. For these other migration schemes, fixation or extinction of defenders was often not reached within the given time frame when *β* was very close to *α*. This may be explained by the fact that under uniform migration, population mixing is strongest as any given deme can be directly reached by individuals. For *simulation series 2D* and *simulation series T*, the final defenders frequency stayed strictly within the boundaries of zero and one for all simulations runs across all parameter settings (contrary to *simulation series U* and *simulation series N*). We conclude that, although the specifics of the population structure do not appear to change which type is favored, they can strongly influence the outcome of the system and lead to prolonged times of coexistence. Thus, the specific spatial structure of a natural population needs to be considered when applying theoretical insights. Across all simulations of *X*, the costly defense trait could only go to fixation (within the simulated time frame of 2000 · *N* generations) if a large number of demes was assumed.

### 4.3 Feedback effects

As Sibly and Curnow (2011) and Berngruber et al. (2013) point out, the success of a costly defense trait depends on the presence of parasites. Thus, benefits of altruistic acts are not constant in the scenario of defense. These fluctuations of effect strengths as well as changing values of relatedness do not need to be considered using the present result. Notably, the current state of host and parasite populations does not enter in the condition for the success of altruism. Instead, the advantage or disadvantage of the defense trait remains fixed over time. Indeed, in most of the simulations, we observed no such explicit negative feedback, and the defense trait typically reaches fixation or extinction. The only exception—for which a negative feedback effect became apparent—was *simulation series g*_*H*_ (Figure 1b). The strong defense of altruists for *β* = 0.0154 led to extinction of parasites in most cases (Section 3.1). If this extinction of parasites occurs before the fixation of altruists, then altruism provides no further benefit and cheaters can take over the population, which may explain why the average final frequency of defenders was slightly lower for *β* = 0.0154 than for any other *β* from 0.005 to 0.012. Thus, the benefit of defense can be detrimental to the evolution of altruism, when parasites go extinct in this finite system. For a biological example see e.g. Duncan et al. (2011), who have shown for the unicellular protist *Paramecium Caudatum* that after removing contact to a parasite, costly resistance is no longer maintained and productivity of the host population slowly increases.

### 4.4 Altruism or only long-term direct fitness advantage of defense?

Deme-level selection can only be effective if there is deme-level variation, and the only source of deme-level variation in our model is genetic drift. The source of genetic drift, however, is the stochasticity of family sizes, which means that that the reason why two individuals sampled from the same deme have an increased probability to be indentical by state (defender or cheater) is that their probability to be identical by decscent is also increased. This allows for a kin selection interpretation of our results because the host indivduals who profit from defense against parasites live in the same deme and tend to be relatives of the defender.

The question is, however, whether the indirect kin selection component is needed at all to explain the advantage of the defense trait. In our model, defense is costly in the sense that defenders have fewer offspring in the next generation than cheaters in the same deme. It could, however, be the case that defense has a direct benefit regarding progeny numbers in later generations due to a reduced parasite load. Even though cheaters in the same deme would also profit from the defense, the defender might in this case have more progeny after a few generations (in the entire population) than it would have if it did turn into a traitor, that is, not apply the defense behavior. We explored this for certain parameter values and found it not to be the case. In fact, traitors in our simulations had on average more progeny after several generations than they had in simulations in which they were actual defenders. Thus, the defense behavior is indeed costly in terms of long-term direct fitness, which justifies calling the defense behavior altruistic, and it seems relevant that among the beneficiaries of the defense are relatives of the defenders besides their own progeny. This being our result for a certain parameter combination does of course not exclude the possibility that for other parameter combinations costly defense behavior can have a long-term direct fitness adavantage. Note that in our simulations in section 3.3 traitors still passed on the defense gene to their offspring, because our objective of these particular simulations was to explore long-term fitness advantage vs. altruism of the defense behavior as such, that is, apart from its inheritance.

### 4.5 Comparison to results obtained with other modeling approaches

Uyenoyama (1979) investigated the evolution of altruism in an island model, assuming a stage of random mixing of the whole population followed by recolonization of islands in each generation. This model goes back to Levene (1953) and has also been utilized by Gillespie (1974). The simplification of assuming a stage of random mixing allows to immediately obtain a one-dimensional diffusion approximation. Uyenoyama (1979) found that for group selection to act, it is not necessary to invoke a mechanism for complete extinction of whole groups. Instead, it is enough if there is maintained variation between the groups, e.g., due to genetic drift or fluctuating environments. Similarly, in our model variation between demes is maintained solely via genetic drift, which is a sufficient force even in large populations, when the processes are analyzed on a large enough time scale. Thus, if some of the other mechanisms to increase and maintain genetic variability (such as mutation, as discussed by Lande, 1976) are introduced in the model, this might provide an even larger benefit for altruism.

Slatkin and Wade (1978) show several mechanisms that are advantageous for the evolution of altruism. Among them are small population sizes per deme, extinction of demes, mutation, and migration in groups with small founding population sizes. None of these are satisfied in the present study, yet, genetic drift produces enough variation between demes in order for deme-level selection to counteract the cost of altruism. Neither the asymptotic analysis of Hutzenthaler et al. (2022) nor our present analysis required the explicit calculation of relatedness values, but it is of course possible to interpret our results from the perspective of inclusive fitness theory. The between-deme variation in the frequency of the defense allele in our model comes from genetic drift and thus from the within-deme relatedness structure. Thus, from the perspective of the causal interpretation of kin selection (Gardner, 2015; Okasha and Martens, 2016) altruists defend their deme because their relatives are overrepresented in the deme.

High amounts of deme-level relatedness also imply, however, that relatives compete for limited space or resources. This competition has been shown to potentially cancel the beneficial effects of cooperation and thus prevent the evolution of altruism (Wilson et al., 1992; Taylor, 1992). However, Alizon and Taylor (2008) show how this effect of competition can be reduced by allowing for empty sites in local populations and varying dispersal rates depending on the size of a deme. Thereby, migration increases with increasing density in a deme and competition can be reduced. We observed a similar effect in the present approach, where larger demes send off more migrants and kin competition can be overcome. However, in our scenario, this effect is obtained with constant migration rates, whereas migration rates are either zero or one (depending on the local population size) in the approach of Alizon and Taylor (2008).

Platt and Bever (2009) review the mechanisms that reduce effects of kin competition and allow for a spread of altruism in structured populations. Apart from allowing for empty sites, an important concept is that of population elasticity. By allowing local carrying capacities to increase under a high density of cooperators, the impact of kin competition can be weakened. In the present model, kin competition is reduced due to population elasticity. Here, a high altruist frequency reduces parasite pressure and thereby increases the potential local host population size.

Débarre et al. (2012) investigated the evolution of altruistic defense traits, in particular suicide upon infection and reduction of transmission, in a host–parasite system with spatial structure modeled as a triangular lattice. They concluded that suicide upon infection with an additional fecundity cost for the ability to detect the parasite can evolve in a structured population if the parasite is sufficiently harmful to the host population. This is in line with the definition of *β* in the present model that captures the parasite-induced selection pressure on hosts as well as the necessity of *κ*_*H*_ being positive, which ensures the population structure in the present model.

## 5 Acknowledgments

For funding we thank the German Science Foundation DFG (grants HU 1889/3-2 and ME 3134/6-2 in Priority Program SPP 1590 “Probabilistc Structures in Evolution”). We thank Georgios Kolyfetis and Solveig Möckel for carrying out preliminary simulations with our simulation program for long-term altruism and Solveig also for proposing the term “traitor” as we now use it in the context of this model.

## Notes

Research supported by the DFG in the Priority Program “Probabilistc Structures in Evolution” (SPP 1590), grants HU 1889/3-2 and ME 3134/6-2.

### Competing Interest Statement

The authors have declared no competing interest.

### Summary of Updates

We found an improved way of matching the finite population-size models to the asymptotic models, which leads to much more accurate predictions whether defenders or non-defenders will reach higher frequencies in host populations. Further we have added additional simulations that show that the defense behavior leads to a decrease in direct fitness, also over more than one generations. Together with the benefit for other host individuals in the same deme, this justifies to call this behavior "altruistic".

