## Supplementary material for "Selection for altruistic defense in structured populations": Online Appendix

### Selection for altruistic defense in structured populations —ONLINE SUPPLEMENT—

March 20, 2023

#### A Diffusion approximation

We now derive the diffusion approximation of the individual-based model defined in Section 2.1. For this, we rescale certain model parameters as in Hutzenthaler et al. (2022) and Hutzenthaler and Metzler (2021) and scale deme population size and (and later time) in units of $N$ individuals (or generations, respectively). Let $\mathcal{D}$ be a finite or countable set that denotes the set of demes and let $m \in [0,\infty)^{\mathcal{D}\times\mathcal{D}}$ such that for every $i \in \mathcal{D}$ it holds that $\sum_{k\in\mathcal{D}} m(k,i) = \sum_{k\in\mathcal{D}} m(i,k) = 1$. Furthermore, let $\lambda$, $K$, $\delta$, $g_H$, $\nu$, $\gamma$, $\eta$, $\rho$, $g_P$, $\alpha$, $\kappa_H$, $\kappa_P \in (0,\infty)$, such that $\rho < \eta$ and let $\sigma \in [0,\infty)^{\mathcal{D}}$ such that $\sum_{i\in\mathcal{D}} \sigma_i < \infty$. Define $E_2 = \left\{x \in [0,\infty)^D \colon \sum_{i\in D} \sigma_i x_i < \infty\right\}$ and let $(\Omega, \mathcal{F}, \mathbb{P})$ be a probability space.

The diffusion approximation will first be performed with unscaled parameters and the scaling will then be introduced for the resulting diffusions to arrive at the appropriate model. For this we denote by $Z_t^N(i) = (A_t^N(i), C_t^N(i), P_t^N(i))$ the group sizes of defenders, cheaters and parasites in units of $N \in \mathbb{N}$ individuals in deme $i \in \mathcal{D}$ at time $t \in [0,\infty)$. (Keep in mind that $.^N$ will be and upper index, not an exponent.) In preparation of the diffusion limit of large deme population sizes, small migration rates and weak selection we parameterize the transition rates of $Z_\cdot^N(.)$ we scale not only within-deme group sizes but also certain parameters with $N$, as defined by the following rates. If we denote for all $k \in \mathcal{D}$ the current state of $Z_t^N(k) \cdot N =$

*corresponding author

$(A_t^N(k)\cdot N, C_t^N(k)\cdot N, P_t^N(k)\cdot N)$ by $(a_k, c_k, p_k)$, a jump of the triple of numbers $(A_t^N(i)\cdot N, C_t^N(i)\cdot N, P_t^N(i)\cdot N)$ to

$$\begin{array}{lll}
(a_i+1, c_i, p_i) & \text{has rate} & a_i(g_H+\lambda) \\
(a_i-1, c_i, p_i) & \text{has rate} & a_i(g_H+\frac{\lambda}{K\cdot N}(a_i+c_i)+\frac{\delta}{N}p_i+\frac{\alpha}{N}) \\
(a_i, c_i+1, p_i) & \text{has rate} & c_i(g_H+\lambda) \\
(a_i, c_i-1, p_i) & \text{has rate} & c_i(g_H+\frac{\lambda}{K\cdot N}(a_i+c_i)+\frac{\delta}{N}p_i) \\
(a_i, c_i, p_i+1) & \text{has rate} & p_i(g_P+\frac{\eta}{N}c_i+\frac{\eta-\rho}{N}a_i) \\
(a_i, c_i, p_i-1) & \text{has rate} & p_i(g_P+\nu+\frac{\gamma}{N}p_i)
\end{array}$$

and for each pair $(i,j)$ of demes with $i \neq j$,

$$\begin{array}{lll}
\multicolumn{3}{l}{(A^N(i)\cdot N, C^N(i)\cdot N, P^N(i)\cdot N) \text{ and } (A^N(j)\cdot N, C^N(j)\cdot N, P^N(j)\cdot N)} \\
\multicolumn{3}{l}{\qquad\text{a simultaneously jump to}} \\
(a_i-1, c_i, p_i) \text{ and } (a_j+1, c_j, p_j) & \text{has rate} & a_i\frac{\kappa_H}{N}m(j,i) \\
(a_i, c_i-1, p_i) \text{ and } (a_j, c_j+1, p_j) & \text{has rate} & c_i\frac{\kappa_H}{N}m(j,i) \\
(a_i, c_i, p_i-1) \text{ and } (a_j, c_j, p_j+1) & \text{has rate} & p_i\frac{\kappa_P}{N}m(j,i).
\end{array} \tag{1}$$

Thus, the per-deme carrying capacity $K$ for a parasite-free host population is measured in units of $N$ individuals, and the host–parasite interaction rates $\delta$, $\eta$, $\rho$ and $\gamma$, which measure the impact of the parasite deme-population size on host individuals and vice versa, are based on population sizes in units of $N$ individuals. For example, $p_i$ is the increase in host death rate caused by the presence of $N$ additional parasite individuals in deme $i$. We interpret the defense selection-cost $\alpha$ and the migration rates $\kappa_H$ and $\kappa_P$ to be per-individual rates measured on the "evolutionary" time scale of $N$ generations. Note that the parameter scalings in section 2.1 of the main text use $N = 1$ and thus refer to $Z^1(.) = (A^1(.), C^1(.), P^1(.))$.

To approximate $Z_t^N(k)$ by a diffusion process, we derive the infinitesimal generator of this process (Ethier and Kurtz, 1986). For this denote by $C(E_2^3, \mathbb{R})$ the set of real-valued continuous functions on $E_2^3$ and by $C_b(E_2^3, \mathbb{R})$ ($C_b^2(E_2^3, \mathbb{R})$) the set of real-valued bounded, continuous functions (that are twice continuously differentiable, with bounded first and second order partial derivatives) on $E_2^3$. Define $\mathrm{Dom}(\mathcal{G}) := \{f \in C_b^2(E_2^3, \mathbb{R})\colon f \text{ depends only on finitely many coordinates}\}$. For $N \in \mathbb{N}$, define the operator $\mathcal{G}_N : \mathrm{Dom}(\mathcal{G}) \to C(E_2^3, \mathbb{R})$, for all $f \in \mathrm{Dom}(\mathcal{G})$ and all $z^N = \left(\frac{a_k}{N}, \frac{c_k}{N}, \frac{p_k}{N}\right)_{k\in\mathcal{D}} \in E_2^3$, such that $\mathbb{P}[Z_0^N = z^N] = 1$, by

$$\mathcal{G}_N f(z^N) = \lim_{t\to 0} \frac{\mathbb{E}[f(Z_t^N) - f(z^N)]}{t}. \tag{2}$$

For all $N \in \mathbb{N}$, for all $f \in \mathrm{Dom}(\mathcal{G})$, and for all $z^N = \left(\frac{a_k}{N}, \frac{c_k}{N}, \frac{p_k}{N}\right)_{k\in\mathcal{D}} \in E_2^3$, such that $\mathbb{P}[Z_0^N = z^N] = 1$, we obtain

$$
\begin{aligned}
\mathcal{G}_N f(z^N) = \lim_{t\to 0} \tfrac{1}{t} \Bigg[ t \sum_{i\in\mathcal{D}} \Bigg( & a_i \Big( g_H + \lambda \Big) f\big( ( \tfrac{a_k + \mathbb{1}_{k=i}}{N}, \tfrac{c_k}{N}, \tfrac{p_k}{N} )_{k\in\mathcal{D}} \big) \\
&+ a_i \Big( g_H + \tfrac{\lambda}{K} \tfrac{a_i + c_i}{N} + \delta \tfrac{p_i}{N} + \tfrac{\alpha}{N} \Big) f\big( ( \tfrac{a_k - \mathbb{1}_{k=i}}{N}, \tfrac{c_k}{N}, \tfrac{p_k}{N} )_{k\in\mathcal{D}} \big) \\
&+ c_i \Big( g_H + \lambda \Big) f\big( ( \tfrac{a_k}{N}, \tfrac{c_k + \mathbb{1}_{k=i}}{N}, \tfrac{p_k}{N} )_{k\in\mathcal{D}} \big) \\
&+ c_i \Big( g_H + \tfrac{\lambda}{K} \tfrac{a_i + c_i}{N} + \delta \tfrac{p_i}{N} \Big) f\big( ( \tfrac{a_k}{N}, \tfrac{c_k - \mathbb{1}_{k=i}}{N}, \tfrac{p_k}{N} )_{k\in\mathcal{D}} \big) \\
&+ p_i \Big( g_P + \eta \tfrac{c_i}{N} + (\eta - \rho) \tfrac{a_i}{N} \Big) f\big( ( \tfrac{a_k}{N}, \tfrac{c_k}{N}, \tfrac{p_k + \mathbb{1}_{k=i}}{N} )_{k\in\mathcal{D}} \big) \\
&+ p_i \Big( g_P + \nu + \gamma \tfrac{p_i}{N} \Big) f\big( ( \tfrac{a_k}{N}, \tfrac{c_k}{N}, \tfrac{p_k - \mathbb{1}_{k=i}}{N} )_{k\in\mathcal{D}} \big) \\
&+ \sum_{j\in\mathcal{D}} a_i \tfrac{\kappa_H}{N} m(j,i) f\big( ( \tfrac{a_k - \mathbb{1}_{k=i} + \mathbb{1}_{k=j}}{N}, \tfrac{c_k}{N}, \tfrac{p_k}{N} )_{k\in\mathcal{D}} \big) \\
&+ \sum_{j\in\mathcal{D}} c_i \tfrac{\kappa_H}{N} m(j,i) f\big( ( \tfrac{a_k}{N}, \tfrac{c_k - \mathbb{1}_{k=i} + \mathbb{1}_{k=j}}{N}, \tfrac{p_k}{N} )_{k\in\mathcal{D}} \big) \\
&+ \sum_{j\in\mathcal{D}} p_i \tfrac{\kappa_P}{N} m(j,i) f\big( ( \tfrac{a_k}{N}, \tfrac{c_k}{N}, \tfrac{p_k - \mathbb{1}_{k=i} + \mathbb{1}_{k=j}}{N} )_{k\in\mathcal{D}} \big) \Bigg) \\
&+ f(z^N) \Big( \mathbb{P}\Big[ Z_t^N = z^N \Big] - 1 \Big) \Bigg].
\end{aligned} \tag{3}
$$

For all $N \in \mathbb{N}$, for all $f \in \mathrm{Dom}(\mathcal{G})$, and for all $z^N = \Big( \frac{a_k}{N}, \frac{c_k}{N}, \frac{p_k}{N} \Big)_{k\in\mathcal{D}} \in E_2^3$, such that $\mathbb{P}[Z_0^N = z^N] = 1$, it follows by Taylor approximation that

$$
\begin{aligned}
\mathcal{G}_N f(z^N) = \sum_{i\in\mathcal{D}} \Bigg\{ & a_i\Big(g_H + \lambda\Big)\Big(\tfrac{d}{da_i} f(z^N)\tfrac{1}{N} + \tfrac{1}{2}\tfrac{d^2}{da_i^2} f(z^N)\tfrac{1}{N^2}\Big) \\
& + a_i\Big(g_H + \tfrac{\lambda}{K}\tfrac{a_i+c_i}{N} + \delta\tfrac{p_i}{N} + \tfrac{\alpha}{N}\Big)\Big( - \tfrac{d}{da_i} f(z^N)\tfrac{1}{N} + \tfrac{1}{2}\tfrac{d^2}{da_i^2} f(z^N)\tfrac{1}{N^2}\Big) \\
& + c_i\Big(g_H + \lambda\Big)\Big(\tfrac{d}{dc_i} f(z^N)\tfrac{1}{N} + \tfrac{1}{2}\tfrac{d^2}{dc_i^2} f(z^N)\tfrac{1}{N^2}\Big) \\
& + c_i\Big(g_H + \tfrac{\lambda}{K}\tfrac{a_i+c_i}{N} + \delta\tfrac{p_i}{N}\Big)\Big( - \tfrac{d}{dc_i} f(z^N)\tfrac{1}{N} + \tfrac{1}{2}\tfrac{d^2}{dc_i^2} f(z^N)\tfrac{1}{N^2}\Big) \\
& + p_i\Big(g_P + \eta\tfrac{c_i}{N} + (\eta-\rho)\tfrac{a_i}{N}\Big)\Big(\tfrac{d}{dp_i} f(z^N)\tfrac{1}{N} + \tfrac{1}{2}\tfrac{d^2}{dp_i^2} f(z^N)\tfrac{1}{N^2}\Big) \\
& + p_i\Big(g_P + \nu + \gamma\tfrac{p_i}{N}\Big)\Big( - \tfrac{d}{dp_i} f(z^N)\tfrac{1}{N} + \tfrac{1}{2}\tfrac{d^2}{dp_i^2} f(z^N)\tfrac{1}{N^2}\Big) \\
& + \sum_{j\in\mathcal{D}} \Bigg[ a_i\tfrac{\kappa_H}{N} m(j,i)\Big( - \tfrac{d}{da_i} f(z^N)\tfrac{1}{N} + \tfrac{d}{da_j} f(z^N)\tfrac{1}{N} + \tfrac{1}{2}\tfrac{d^2}{da_i da_j} f(z^N)\tfrac{1}{N^2}\Big) \\
& + c_i\tfrac{\kappa_H}{N} m(j,i)\Big( - \tfrac{d}{dc_i} f(z^N)\tfrac{1}{N} + \tfrac{d}{dc_j} f(z^N)\tfrac{1}{N} + \tfrac{1}{2}\tfrac{d^2}{dc_i dc_j} f(z^N)\tfrac{1}{N^2}\Big) \\
& + p_i\tfrac{\kappa_P}{N} m(j,i)\Big( - \tfrac{d}{dp_i} f(z^N)\tfrac{1}{N} + \tfrac{d}{dp_j} f(z^N)\tfrac{1}{N} + \tfrac{1}{2}\tfrac{d^2}{dp_i dp_j} f(z^N)\tfrac{1}{N^2}\Big) \Bigg] \\
& + o(\tfrac{1}{N^2}) \Bigg\}.
\end{aligned}
\tag{4}
$$

To simplify the migration terms we use for all $i \in \mathcal{D}$ that $\sum_{k\in\mathcal{D}} m(i,k) = \sum_{k\in\mathcal{D}} m(k,i) = 1$. By rearranging terms for all $N \in \mathbb{N}$, for all $f \in \mathrm{Dom}(\mathcal{G})$, and for all $z^N =$

$\left(\frac{a_k}{N}, \frac{c_k}{N}, \frac{p_k}{N}\right)_{k\in\mathcal{D}} \in E_2^3$, such that $\mathbb{P}[Z_0^N = z^N] = 1$, it follows that

$$\begin{aligned}
&\sum_{i\in\mathcal{D}}\sum_{j\in\mathcal{D}}\Bigg[a_i\tfrac{\kappa_H}{N}m(j,i)\Big(-\tfrac{d}{da_i}f(z^N)\tfrac{1}{N}+\tfrac{d}{da_j}f(z^N)\tfrac{1}{N}+\tfrac{1}{2}\tfrac{d^2}{da_i da_j}f(z^N)\tfrac{1}{N^2}\Big)\\
&\quad+c_i\tfrac{\kappa_H}{N}m(j,i)\Big(-\tfrac{d}{dc_i}f(z^N)\tfrac{1}{N}+\tfrac{d}{dc_j}f(z^N)\tfrac{1}{N}+\tfrac{1}{2}\tfrac{d^2}{dc_i dc_j}f(z^N)\tfrac{1}{N^2}\Big)\\
&\quad+p_i\tfrac{\kappa_P}{N}m(j,i)\Big(-\tfrac{d}{dp_i}f(z^N)\tfrac{1}{N}+\tfrac{d}{dp_j}f(z^N)\tfrac{1}{N}+\tfrac{1}{2}\tfrac{d^2}{dp_i dp_j}f(z^N)\tfrac{1}{N^2}\Big)\Bigg]\\
&=\sum_{i\in\mathcal{D}}\Bigg[-\tfrac{a_i}{N}\tfrac{\kappa_H}{N}\tfrac{d}{da_i}f(z^N)+\sum_{j\in\mathcal{D}}\tfrac{a_j}{N}\tfrac{\kappa_H}{N}m(i,j)\tfrac{d}{da_i}f(z^N)-\tfrac{c_i}{N}\tfrac{\kappa_H}{N}\tfrac{d}{dc_i}f(z^N)\\
&\quad+\sum_{j\in\mathcal{D}}\tfrac{c_j}{N}\tfrac{\kappa_H}{N}m(i,j)\tfrac{d}{dc_i}f(z^N)-\tfrac{p_i}{N}\tfrac{\kappa_P}{N}\tfrac{d}{dp_i}f(z^N)+\sum_{j\in\mathcal{D}}\tfrac{p_j}{N}\tfrac{\kappa_P}{N}m(i,j)\tfrac{d}{dp_i}f(z^N)\\
&\quad+\sum_{j\in\mathcal{D}}m(j,i)\cdot o(\tfrac{1}{N^2})\Bigg]\\
&=\sum_{i\in\mathcal{D}}\sum_{j\in\mathcal{D}}\Bigg[\tfrac{\kappa_H}{N}m(i,j)\Big(\tfrac{a_j}{N}-\tfrac{a_i}{N}\Big)\tfrac{d}{da_i}f(z^N)+\tfrac{\kappa_H}{N}m(i,j)\Big(\tfrac{c_j}{N}-\tfrac{c_i}{N}\Big)\tfrac{d}{dc_i}f(z^N)\\
&\quad+\tfrac{\kappa_P}{N}m(i,j)\Big(\tfrac{p_j}{N}-\tfrac{p_i}{N}\Big)\tfrac{d}{dp_i}f(z^N)\Bigg]+\sum_{i\in\mathcal{D}}o(\tfrac{1}{N^2}).
\end{aligned} \tag{5}$$

With this, we obtain for all $N \in \mathbb{N}$, all $f \in \mathrm{Dom}(\mathcal{G})$, and $z^N = \left(\frac{a_k}{N}, \frac{c_k}{N}, \frac{p_k}{N}\right)_{k\in\mathcal{D}} \in$

$E_2^3$ with $\mathbb{P}[Z_0^N = z^N] = 1$ the approximation

$$\mathcal{G}_N f(z^N) = \sum_{i\in\mathcal{D}} \Bigg\{ \tfrac{a_i}{N}\Big(\lambda - \tfrac{\lambda}{K}\tfrac{a_i+c_i}{N} - \delta\tfrac{p_i}{N} - \tfrac{\alpha}{N}\Big)\tfrac{d}{da_i}f(z^N)$$
$$+ \tfrac{a_i}{2N}(2g_H + \lambda + \tfrac{\lambda}{K}\tfrac{a_i+c_i}{N} + \delta\tfrac{p_i}{N} + \tfrac{\alpha}{N}\Big)\tfrac{d^2}{da_i^2}f(z^N)\tfrac{1}{N}$$
$$+ \tfrac{c_i}{N}\Big(\lambda - \tfrac{\lambda}{K}\tfrac{a_i+c_i}{N} - \delta\tfrac{p_i}{N}\Big)\tfrac{d}{dc_i}f(z^N)$$
$$+ \tfrac{c_i}{2N}(2g_H + \lambda + \tfrac{\lambda}{K}\tfrac{a_i+c_i}{N} + \delta\tfrac{p_i}{N})\tfrac{d^2}{dc_i^2}f(z^N)\tfrac{1}{N}$$
$$+ \tfrac{p_i}{N}\Big(-\nu - \gamma\tfrac{p_i}{N} + \eta\tfrac{c_i}{N} + (\eta-\rho)\tfrac{a_i}{N}\Big)\tfrac{d}{dp_i}f(z^N)$$
$$+ \tfrac{p_i}{2N}(2g_P + \nu + (\eta-\rho)\tfrac{a_i}{N} + \eta\tfrac{c_i}{N} + \gamma\tfrac{p_i}{N})\tfrac{d^2}{dp_i^2}f(z^N)\tfrac{1}{N}$$
$$+ \sum_{j\in\mathcal{D}} \tfrac{\kappa_H}{N} m(i,j)\Big(\tfrac{a_j}{N} - \tfrac{a_i}{N}\Big)\tfrac{d}{da_i}f(z^N) + \sum_{j\in\mathcal{D}} \tfrac{\kappa_H}{N} m(i,j)\Big(\tfrac{c_j}{N} - \tfrac{c_i}{N}\Big)\tfrac{d}{dc_i}f(z^N)$$
$$+ \sum_{j\in\mathcal{D}} \tfrac{\kappa_P}{N} m(i,j)\Big(\tfrac{p_j}{N} - \tfrac{p_i}{N}\Big)\tfrac{d}{dp_i}f(z^N) + o(\tfrac{1}{N^2})\Bigg\}.$$
$$= \sum_{i\in\mathcal{D}} \Bigg\{ \Bigg[\tfrac{\kappa_H}{N}\sum_{j\in\mathcal{D}} m(i,j)\Big(\tfrac{a_j}{N} - \tfrac{a_i}{N}\Big) + \tfrac{a_i}{N}\Big(\lambda - \tfrac{\lambda}{K}\tfrac{a_i+c_i}{N} - \delta\tfrac{p_i}{N} - \tfrac{\alpha}{N}\Big)\Bigg]\tfrac{d}{da_i}f(z^N)$$
$$+ \Bigg[\tfrac{\kappa_H}{N}\sum_{j\in\mathcal{D}} m(i,j)\Big(\tfrac{c_j}{N} - \tfrac{c_i}{N}\Big) + \tfrac{c_i}{N}\Big(\lambda - \tfrac{\lambda}{K}\tfrac{a_i+c_i}{N} - \delta\tfrac{p_i}{N}\Big)\Bigg]\tfrac{d}{dc_i}f(z^N)$$
$$+ \Bigg[\tfrac{\kappa_P}{N}\sum_{j\in\mathcal{D}} m(i,j)\Big(\tfrac{p_j}{N} - \tfrac{p_i}{N}\Big) + \tfrac{p_i}{N}\Big(-\nu - \gamma\tfrac{p_i}{N} + \eta\tfrac{c_i}{N} + (\eta-\rho)\tfrac{a_i}{N}\Big)\Bigg]\tfrac{d}{dp_i}f(z^N)$$
$$+ \tfrac{a_i}{2N}(2g_H + \lambda + \tfrac{\lambda}{K}\tfrac{a_i+c_i}{N} + \delta\tfrac{p_i}{N} + \tfrac{\alpha}{N})\tfrac{d^2}{da_i^2}f(z^N)\tfrac{1}{N}$$
$$+ \tfrac{c_i}{2N}(2g_H + \lambda + \tfrac{\lambda}{K}\tfrac{a_i+c_i}{N} + \delta\tfrac{p_i}{N})\tfrac{d^2}{dc_i^2}f(z^N)\tfrac{1}{N}$$
$$+ \tfrac{p_i}{2N}(2g_P + \nu + (\eta-\rho)\tfrac{a_i}{N} + \eta\tfrac{c_i}{N} + \gamma\tfrac{p_i}{N})\tfrac{d^2}{dp_i^2}f(z^N)\tfrac{1}{N}$$
$$+ o(\tfrac{1}{N^2})\Bigg\}. \tag{6}$$

With this approximation of the generator we obtain a diffusion approximation (Ethier and Kurtz, 1986) of $Z_t^N(i)_{i\in\mathcal{D}} = (A_t^N(i), C_t^N(i), P_t^N(i))_{i\in\mathcal{D}}$ for large $N$ of the following form. With independent Brownian motions $W^{A,N}(i), W^{C,N}(i), W^{P,N}(i)\colon [0,\infty)\times\Omega \to \mathbb{R}$, $i \in \mathcal{D}$, with continuous sample paths, $A^N(i)$, $C^N(i)$, $P^N(i)\colon [0,\infty)\times\mathcal{D}\times\Omega \to [0,\infty)$, are adapted processes with continuous sample paths that for all $i \in \mathcal{D}$ and all $t \in [0,\infty)$ satisfy $\mathbb{P}$-a.s.

$$
\begin{aligned}
A_t^N(i) &= A_0^N(i) + \int_0^t \Bigg( \tfrac{\kappa_H}{N} \sum_{j\in\mathcal{D}} m(i,j)\left(A_s^N(j) - A_s^N(i)\right) \\
&\quad + A_s^N(i)\Big[\lambda\left(1 - \tfrac{A_s^N(i)+C_s^N(i)}{K}\right) - \delta P_s^N(i) - \tfrac{\alpha}{N}\Big]\Bigg)\, ds \\
&\quad + \int_0^t \sqrt{\tfrac{1}{N}\left(2g_H + \lambda + \tfrac{\lambda\cdot(A_s^N(i)+C_s^N(i))}{K} + \delta\cdot P_s^N(i) + \frac{\alpha}{N}\right) A_s^N(i)}\, dW_s^{A,N}(i), \\
C_t^N(i) &= C_0^N(i) + \int_0^t \Bigg( \tfrac{\kappa_H}{N} \sum_{j\in\mathcal{D}} m(i,j)\left(C_s^N(j) - C_s^N(i)\right) \\
&\quad + C_s^N(i)\Big[\lambda\left(1 - \tfrac{A_s^N(i)+C_s^N(i)}{K}\right) - \delta P_s^N(i)\Big]\Bigg)\, ds \\
&\quad + \int_0^t \sqrt{\tfrac{1}{N}\left(2g_H + \lambda + \tfrac{\lambda\cdot(A_s^N(i)+C_s^N(i))}{K} + \delta\cdot P_s^N(i)\right) C_s^N(i)}\, dW_s^{C,N}(i), \\
P_t^N(i) &= P_0^N(i) + \int_0^t \Bigg( \tfrac{\kappa_P}{N} \sum_{j\in\mathcal{D}} m(i,j)\left(P_s^N(j) - P_s^N(i)\right) + P_s^N(i)\Big[-\nu - \gamma P_s^N(i) \\
&\quad + \eta C_s^N(i) + (\eta-\rho)A_s^N(i)\Big]\Bigg)\, ds \\
&\quad + \int_0^t \sqrt{\tfrac{1}{N}\left(2g_P + \nu + (\eta-\rho)A_s^N(i) + \eta C_s^N(i) + \gamma P_s^N(i)\right) P_s^N(i)}\, dW_s^{P,N}(i).
\end{aligned}
\tag{7}
$$

We simplify this diffusion model for the case that the effect $\rho$ of defense and its cost $\alpha/N$ are small and that the system is not too far from the Lotka–Volterra equilibrium in each deme, such that the host death rate $g_H + \frac{\lambda\cdot(A_s^N(i)+C_s^N(i))}{K} + \delta\cdot P_s^N(i)$ (with or without the small summand $\frac{\alpha}{N}$) is approximately the host birth rate $g_H + \lambda$. If we neglect the effect of defense, the Lotka–Volterra equilibrium population size in a deme is $\frac{\lambda\cdot(K\eta-\nu)}{\delta\eta K+\gamma\lambda}$. With this population size, the death rate $g_H + \nu + \delta\cdot P_s^N(i)$ of parasites is $g_H + \nu + \gamma\frac{\lambda\cdot(K\eta-\nu)}{\delta\eta K+\gamma\lambda}$, and if the system is close to equilibrium, the total parasite birth rate $g_P + \nu + (\eta-\rho)A_s^N(i) + \eta C_s^N(i)$ must be a similar value. Applying these heuristic equilibrium-based approximations to the diffusion terms (and only there!) we obtain the following diffusion model with slightly simpler diffusion term:

$$
\begin{aligned}
A_t^N(i) &= A_0^N(i) + \int_0^t \Bigg( \tfrac{\kappa_H}{N} \sum_{j\in\mathcal{D}} m(i,j)\left(A_s^N(j) - A_s^N(i)\right) \\
&\quad + A_s^N(i)\Big[\lambda\left(1 - \tfrac{A_s^N(i)+C_s^N(i)}{K}\right) - \delta P_s^N(i) - \tfrac{\alpha}{N}\Big]\Bigg)\,ds \\
&\quad + \int_0^t \sqrt{\tfrac{1}{N}\left(2g_H + 2\lambda\right) A_s^N(i)}\, dW_s^{A,N}(i), \\
C_t^N(i) &= C_0^N(i) + \int_0^t \Bigg( \tfrac{\kappa_H}{N} \sum_{j\in\mathcal{D}} m(i,j)\left(C_s^N(j) - C_s^N(i)\right) \\
&\quad + C_s^N(i)\Big[\lambda\left(1 - \tfrac{A_s^N(i)+C_s^N(i)}{K}\right) - \delta P_s^N(i)\Big]\Bigg)\,ds \\
&\quad + \int_0^t \sqrt{\tfrac{1}{N}\left(2g_H + 2\lambda\right) C_s^N(i)}\, dW_s^{C,N}(i), \\
P_t^N(i) &= P_0^N(i) + \int_0^t \Bigg( \tfrac{\kappa_P}{N} \sum_{j\in\mathcal{D}} m(i,j)\left(P_s^N(j) - P_s^N(i)\right) \\
&\quad + P_s^N(i)\Big[-\nu - \gamma P_s^N(i) + \eta C_s^N(i) + (\eta-\rho) A_s^N(i)\Big]\Bigg)\,ds \\
&\quad + \int_0^t \sqrt{\tfrac{2}{N}\left(g_P + \nu + \tfrac{\lambda\gamma\cdot(K\eta-\nu)}{\lambda\gamma+\delta K\nu}\right) P_s^N(i)}\, dW_s^{P,N}(i).
\end{aligned}
\tag{8}
$$

Equation system (8) specifies the model for which Hutzenthaler et al. (2022) showed that the asymptotic behavior of the defense allele frequencies $\left(\frac{A_t^N(i)}{A_t^N(i)+C_t^N(i)}\right)_{i\in\mathcal{D}}$ in the limit of large demes and evolutionary time scaling is given by diffusion equation system (1) (see main text).

| Par. | Description | $\varepsilon$ |
|---|---|---|
| $\beta$ | random fluctuation (“benefit”) | 0.01 to 0.09 |
| $\alpha$ | altruist disadvantage | 0.05 |
| $\kappa$ | migration rate | 0.1 |
| $a$ | see Section 2.3 | 2 |
| $D$ | number of demes | 500 |
| $x_0$ | initial altruist frequency per deme | 0.5 |
| $T$ | time horizon | 2000 |
| $dt$ | time step size of discretization | $10^{-5}$ |

Table 1: Parameters of diffusion simulations for *simulation series* $\varepsilon$

### B Simulation series $\varepsilon$

The discretization in the simulation procedure in Section 2.3 may introduce a bias away from the boundaries if $X_i$ attains values very close to zero or one. This is due to the fact that near zero or one, steps in direction of the boundaries cannot exceed the boundaries, while steps away from the boundaries are taken in full size. To compensate for this, values below $\varepsilon$ or above $1 - \varepsilon$, are set to the corresponding boundary values of zero or one. In *simulation series* $\varepsilon$, a suitable cutoff value was determined. Here, suitable means large enough such that the bias does not affect the outcome and small enough such that immigration can still drive demes away from fixation or extinction. Note, however, that if the process is simulated for a very long time, this effect may be overcome by chance. For each parameter set, 25 simulations were performed, with parameters given in Table 1. Since in *simulation series* $\varepsilon$, $\alpha$ is set to $0.05$, the predicted value is zero for $\beta < 0.05$ and one for $\beta > 0.05$. Final altruist frequencies for all runs are shown in Figure 1. Points are spread out horizontally. The actual values of $\beta$ are $0.01$, $0.03$, $0.07$, and $0.09$. Extinction or fixation of altruists occurred in all runs for $\varepsilon = 10^{-7}$, in 97 of the 100 runs for $\varepsilon = 10^{-6}$, in the 25 runs with $\beta = 0.01$ but never with other values of $\beta$ for $\varepsilon = 10^{-8}$, and in none of the runs for $\varepsilon = 0$.

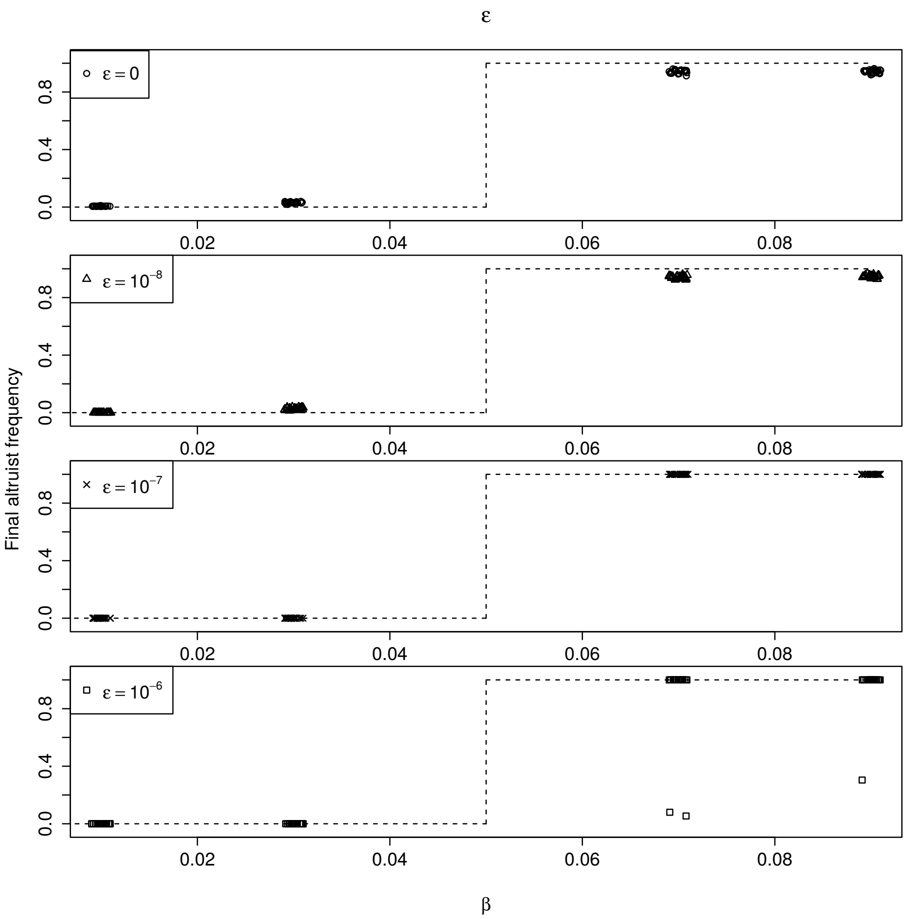


Figure 1: Final altruist frequencies from *simulation series* $\varepsilon$. Points are spread out horizontally. Dashed gray lines indicate the theoretical predictions.

### C Simulations for altruist frequency in large populations

In this section, further results from the simulation study are presented. Details are given in Section 3.2.

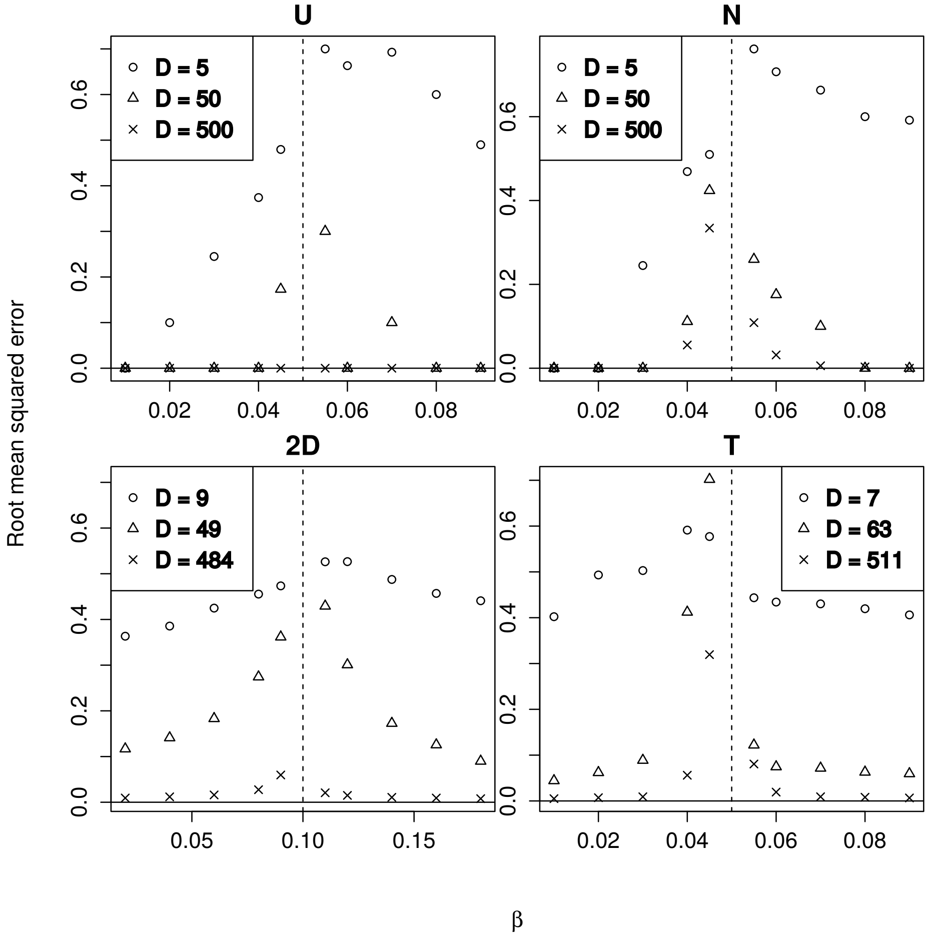


Figure 2: Root mean squared errors from 100 simulation runs with uniform migration (U), nearest-neighbor migration (N), two-dimensional nearest-neighbor migration (2D), and migration along edges of a binary tree (T). Dashed gray lines indicate the respective values of $\alpha$. The simulations with $\beta = \alpha$ were excluded from the computation of root mean squared errors.

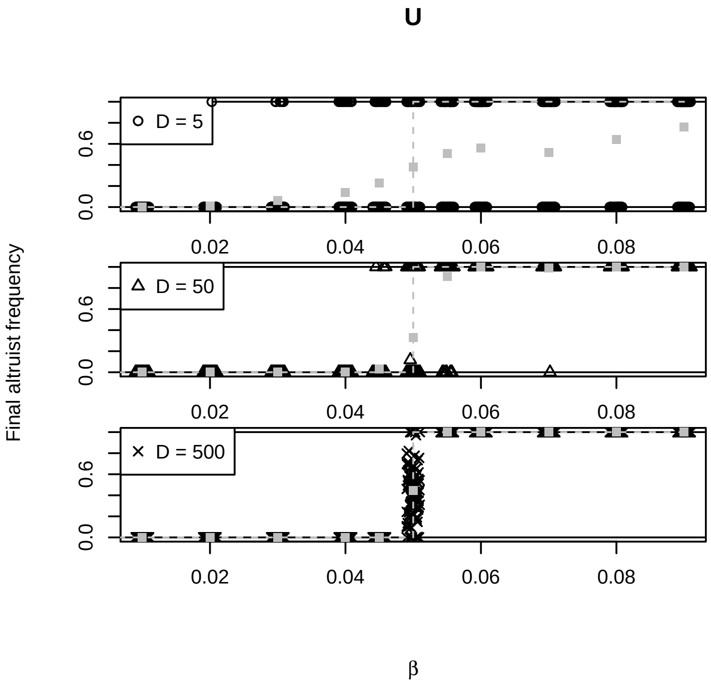


Figure 3: Final altruist frequencies from *simulation series U*. Dashed gray lines indicate the theoretical predictions. Each point indicates the final altruist frequency of one simulation run. Points are spread out horizontally. The actual values of $\beta$ are $0.01$, $0.02$, $0.03$, $0.04$, $0.045$, $0.05$, $0.055$, $0.06$, $0.07$, $0.08$, and $0.09$.

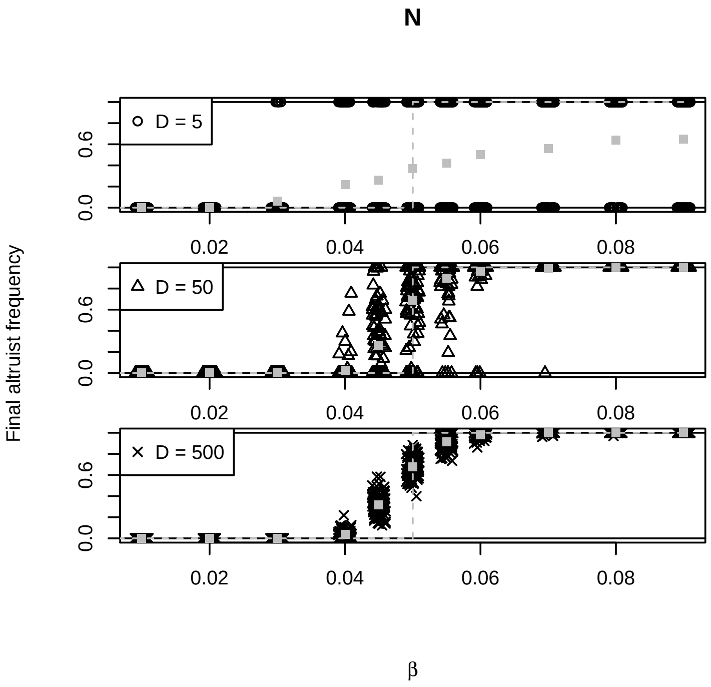


Figure 4: Final altruist frequencies from *simulation series N*. Dashed gray lines indicate the theoretical predictions. Each point indicates the final altruist frequency of one simulation run. Points are spread out horizontally. The actual values of $\beta$ are 0.01, 0.02, 0.03, 0.04, 0.045, 0.05, 0.055, 0.06, 0.07, 0.08, and 0.09.

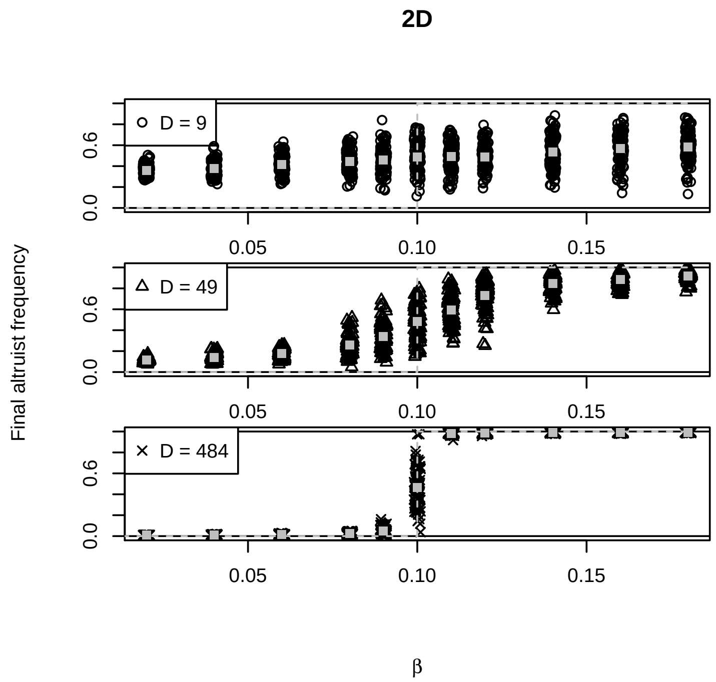


Figure 5: Final altruist frequencies from *simulation series 2D*. Dashed gray lines indicate the theoretical predictions. Each point indicates the final altruist frequency of one simulation run. Points are spread out horizontally. The actual values of $\beta$ are $0.02$, $0.04$, $0.06$, $0.08$, $0.09$, $0.1$, $0.11$, $0.12$, $0.14$, $0.16$, and $0.18$.

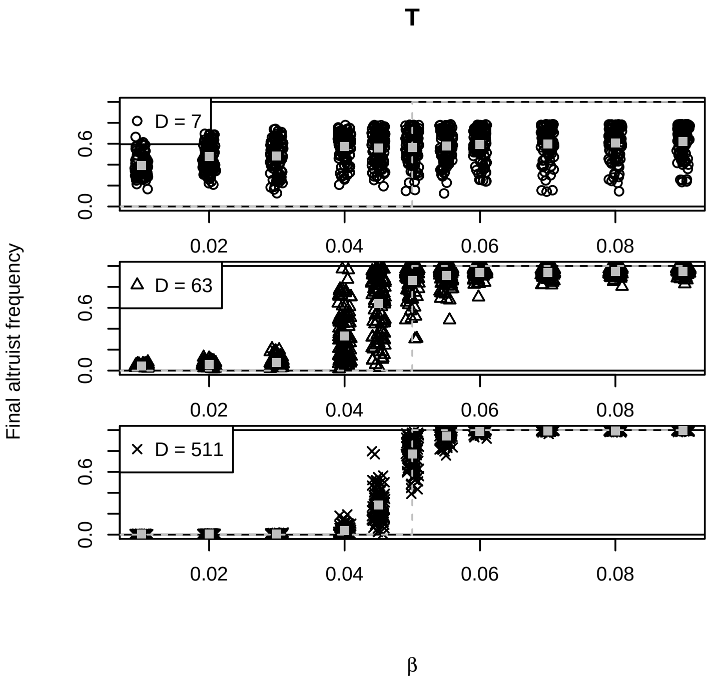


Figure 6: Final altruist frequencies from *simulation series T*. Points are spread out horizontally. Dashed gray lines indicate the theoretical predictions. Each point indicates the final altruist frequency of one simulation run. Points are spread out horizontally. The actual values of $\beta$ are 0.01, 0.02, 0.03, 0.04, 0.045, 0.05, 0.055, 0.06, 0.07, 0.08, and 0.09.

### D $\tau$-leap simulations for comparison with individual-based simulation

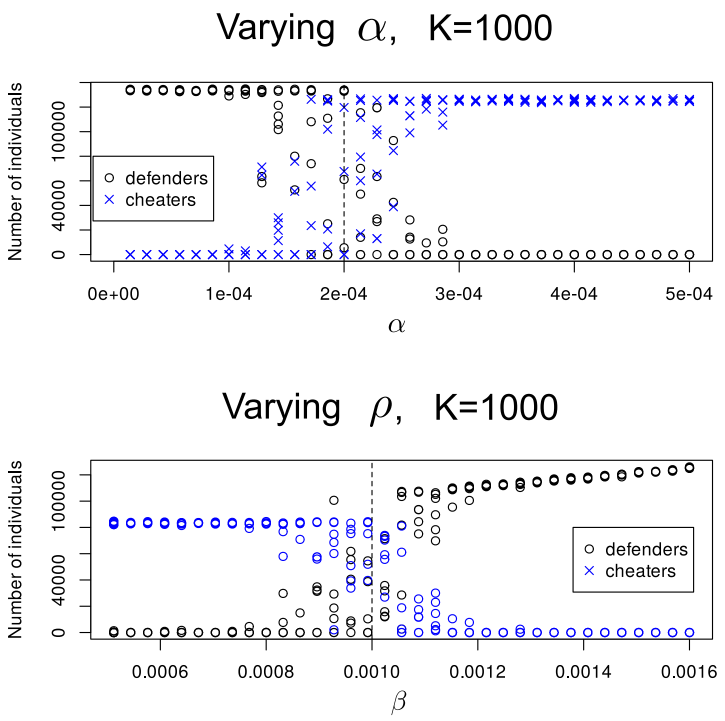


Figure 7: Numbers of defenders and cheaters after phase 1 of simulations as specified in section 2.4 of the main text. Dashed lines show the threshold accoring to the asymptotic diffusion model. Top: parameters like in series $\alpha$ with $\rho = 1.24 \cdot 10^{-4}$, bottom: parameters like in series $g_H$ and $k_H$
